# Megadomains and superloops form dynamically but are dispensable for X-chromosome inactivation and gene escape

**DOI:** 10.1101/364893

**Authors:** John E. Froberg, Stefan F. Pinter, Andrea J. Kriz, Teddy Jégu, Jeannie T. Lee

## Abstract

The mammalian inactive X-chromosome (Xi) is structurally distinct from all other chromosomes and serves as a model for how the 3D genome is organized. The Xi shows weakened topologically associated domains and is instead organized into megadomains and superloops directed by the noncoding loci, *Dxz4* and *Firre*. Their functional significance is presently unclear, though one study suggests that they permit Xi genes to escape silencing. Here, we find that megadomains do not precede Xist expression or Xi gene silencing. Deleting *Dxz4* disrupts megadomain formation, whereas deleting *Firre* weakens intra-megadomain interactions. Surprisingly, however, deleting *Dxz4* and *Firre* has no impact on Xi silencing and gene escape. Nor does it affect Xi nuclear localization, stability, or H3K27 methylation. Additionally, ectopic integration of *Dxz4* and *Xist* is not sufficient to form megadomains on autosomes, further uncoupling megadomain formation from chromosomal silencing. We conclude that *Dxz4* and megadomains are dispensable for Xi silencing and escape from X-inactivation.

## INTRODUCTION

A longstanding principle in gene regulation invokes significance of higher order chromosome structures and specificity of 3D interactions between distant genetic elements. Advances in genomics have provided new opportunities to probe chromosome architecture and resulted in discovery of three types of long-range intrachromosomal interactions. First, “topologically associating domains” (TADs) define continuous regions with extensive cis-contacts ^1^. TADs are usually observed at length scales from 10^4^-10^6^ bp^2^, depend on cohesins^3-5^,and are generally flanked by convergent CTCF sites at TAD borders^1,6-8^. TADs are visible as squares along the diagonal of Hi-C contact heat maps. Second, “loops” define enhanced contacts between pairs of distant loci that interact via CTCF. Loops can exist within or between TADs, and are visible as strong “dots” within a TAD square or between separate TADs in Hi-C contact maps^1^. Third, “compartments” transcend TADs and loops and exist as orthogonal structures formed by interactions between chromatin of similar epigenetic states. A-compartments harbor discontinuous chromosomal regions enriched for active genes, whereas B-compartments harbor discontiguous regions enriched for repressed genes^1,3,4,9-11^. A/B compartments are visualized in Hi-C correlation maps by alternating “plaid” patterns of strong and weak interactions. Rapid depletion of CTCF^10^ or cohesin^3^ leads to genome-wide loss of TADs and loops, more pronounced A/B compartments (in the case of cohesin depletion) and only modestly affects transcription in the short term^3-5,12^. Thus, while loops are thought to be important for long-range gene regulation (such as enhancer-promoter interactions), the functional organization into TADs and compartments is presently less well understood.

Recent conformational studies of the inactive X (Xi) has provided new insight into 3D chromosomal structure-function relationships^13,14^. X-chromosome inactivation (XCI) occurs in female cells as part of a dosage compensation mechanism that equalizes dosage of X-linked genes between males and females^15-17^. Chromosome conformation capture studies have demonstrated that, whereas the active X (Xa) resembles autosomes in having defined TADs, loops, and compartments, the Xi adopts a distinct structure seen on no other mammalian chromosome^1,18-22^. ChIP-seq studies have shown that binding of architectural proteins including CTCF^20,23^ and cohesins^20^ are relatively depleted on the Xi, providing a mechanistic explanation for the attenuation of TADs. The Xi also lacks the characteristic separation between transcriptionally active A compartments and silent B compartments. Instead, during XCI, the Xi is partitioned into transitional Xist-rich S1 and Xist-poor S2 compartments, which are later merged into a single compartment by the non-canonical SMC protein, SMCHD1^24^. The merging of S1/S2 structure has physiological consequence, as ablating SMCHD1 precludes this fusion and leads to failure of silencing of >40% of genes on the Xi ^24^. Thus, on the Xi, compartmentalization appears to play an important role in gene silencing.

On the other hand, the significance of domains on the Xi is under debate. Studies have shown that the Xi folds into two “megadomains” separated by a non-coding locus bearing tandem repeats known as “*Dxz4*” ^20-22,25^. In humans, *DXZ4* is heterochromatinized and methylated on the Xa but is euchromatic and unmethylated on the Xi, where it binds CTCF^26,27^. Murine *Dxz4* is not well-conserved at the sequence level, but the syntenic region harbors a unique tandem repeat harboring strong CTCF binding sites^28^. In both mouse and human, *Dxz4/DXZ4* resides at the strong border between the two megadomains of the Xi and binds CTCF and cohesin in an allele-specific manner (Fig. S1). Deleting the *Dxz4/DXZ4* region in both species results in loss of megadomains and increased frequency of interaction across the border^22,25,29^. Despite clear disruption of the Xi super-structure, there is presently no agreement regarding functional consequences. One group reported loss of ability of “escapees” to avoid silencing on the Xi^22^. Changes in repressive chromatin marks and accessibility have also been reported in the mouse^22,29^. Still others reported minimal effects, or even opposite effects, such as a partial loss of Xi heterochromatin in human cells^25^. Thus, there exists major disagreement as to whether the *Dxz4* region and megadomains enable or oppose silencing.

Additionally, the Xi is characterized by a network of extremely long-range loops termed “superloops”^1^ and the importance of these structures is also unknown. Superlooping occurs between *Xist, DXZ4,* and another tandem repeat element on the Xi called *FIRRE*^30,31^, another CTCF-bound noncoding locus (Fig. S1)^1,25,27^. Far longer than almost all other contacts in mammalian genomes, the loops between *Dxz4* and *Firre* extrude up to 25 Mb of DNA, a scale typically seen only in perturbed states, such as between super-enhancers of cohesin-depleted cells^3^. One study suggests that Firre RNA may direct Xist to the perinucleolar space and influence H3K27me3 deposition on the X^32^. However, despite the fact that the *Firre* locus falls at the border between two TADs and contains many CTCF binding sites, a recent study found that *Firre* is neither necessary nor sufficient to form borders between TADs, though it is required for superlooping with *Dxz4^33^*. Here we combine genetic, epigenomic, and cell biological methods and study the impact of large-scale 3D structures on Xi biology.

## RESULTS

### Megadomains appear after Xist expression but not before Xi gene silencing

It is presently unknown how the formation of Xi megadomains relates to the timeline of XCI. To assess whether megadomains precede or follow XCI, we performed allele-specific Hi-C in female mouse embryonic stem cells (ES), which model different steps of XCI when they are induced to differentiate in culture. We examined timepoints day 0 (before XCI), day 3 (early XCI), day 7 (mid-XCI), and day 10 (late-XCI) in the mESC line, *Tsix^TST^/+*, and compared the megadomain timeline to the time course of Xist upregulation and Xi silencing across the differentiating population in two biological replicates. Allelic analysis was made possible in *Tsix^TST^/+* in two ways. First, it carries one X-chromosome of *M. castaneus* (cas) origin and one of *M. musculus* 129 origin (mus)^34,35^, the combination of which enabled employment of >600,000 polymorphisms to distinguish alleles. Second, the cell line carries a stop-mutation in the mus *Tsix* allele^34^ that ensures that the mus X-chromosome is chosen as the Xi in >95% of cells^35-37^.

For Hi-C analysis, we sequenced to a depth of 25-50 million reads, as megadomains are large (>70 Mb), prominent structures and can be sensitively detected at a resolution of 2.5 megabases (Mb). Allele-specific Hi-C contact maps and corresponding Pearson correlation heatmaps showed that, as expected, megadomains did not appear on the Xa during any stage of differentiation (Fig. S2a,b). Focusing in on the Xi, we observed that, in pre-XCI cells (day 0) and in cells undergoing XCI (day 3), X^mus^ resembled X^cas^ (the Xa) in also lacking detectable megadomains (Fig. 1a,b). RNA FISH analysis of these timepoints showed that 30-60% of cells showed robust Xist RNA clouds by day 3 (Fig. 1c,d). Allele-specific RNA-seq analysis also showed robust upregulation of Xist starting on day 3 and continuing throughout differentiation (Fig. 1e, Fig S2c,d). Importantly, Xist was upregulated almost exclusively from X^mus^ as expected, consistent with the *Tsix^TST^* allele carried in cis^34^. [*Note: The nonrandom pattern at day 3 agrees with Tsix being a primary determinant of allelic choice^38^ rather than being a secondary selection mechanism following a stochastic choice process^39,40^. A small fraction of reads coming from* X^cas^ *is likely to be artifactual, as virtually all the* X^cas^ *reads felll into one peak near the 5’ end of Xist, rather than being distributed across the entire gene body (Fig. 1e). This peak fell within a repetitive region of Xist (repeat A) and contained only one SNP (<u>rs225651233</u>) — a 129 G -> Cast/EiJ T variant falling within a low complexity 24 bp poly-T tract. Thus, the* X^cas^ *reads are likely to be from an improperly-defined SNP.*] Despite highly skewed Xist upregulation, allele-specific RNAseq analysis showed that X-linked gene expression remained relatively unskewed (Fig. 1f), implying that de novo silencing or turnover of preexisting mRNA lagged behind Xist upregulation. A Hi-C “mixing” experiment indicated that our allelic Hi-C assay could detect megadomains when present in 25% of cells (Fig. 1h,i). Thus, the fact that megadomains were not readily visible on day 3 suggests that <25% of cells harbored them. Given that Xist spreading had taken place in 30-60% of cells and little silencing had taken place at this time point, megadomains were unlikely to have preceded Xist spreading and gene silencing.

**Figure 1:**
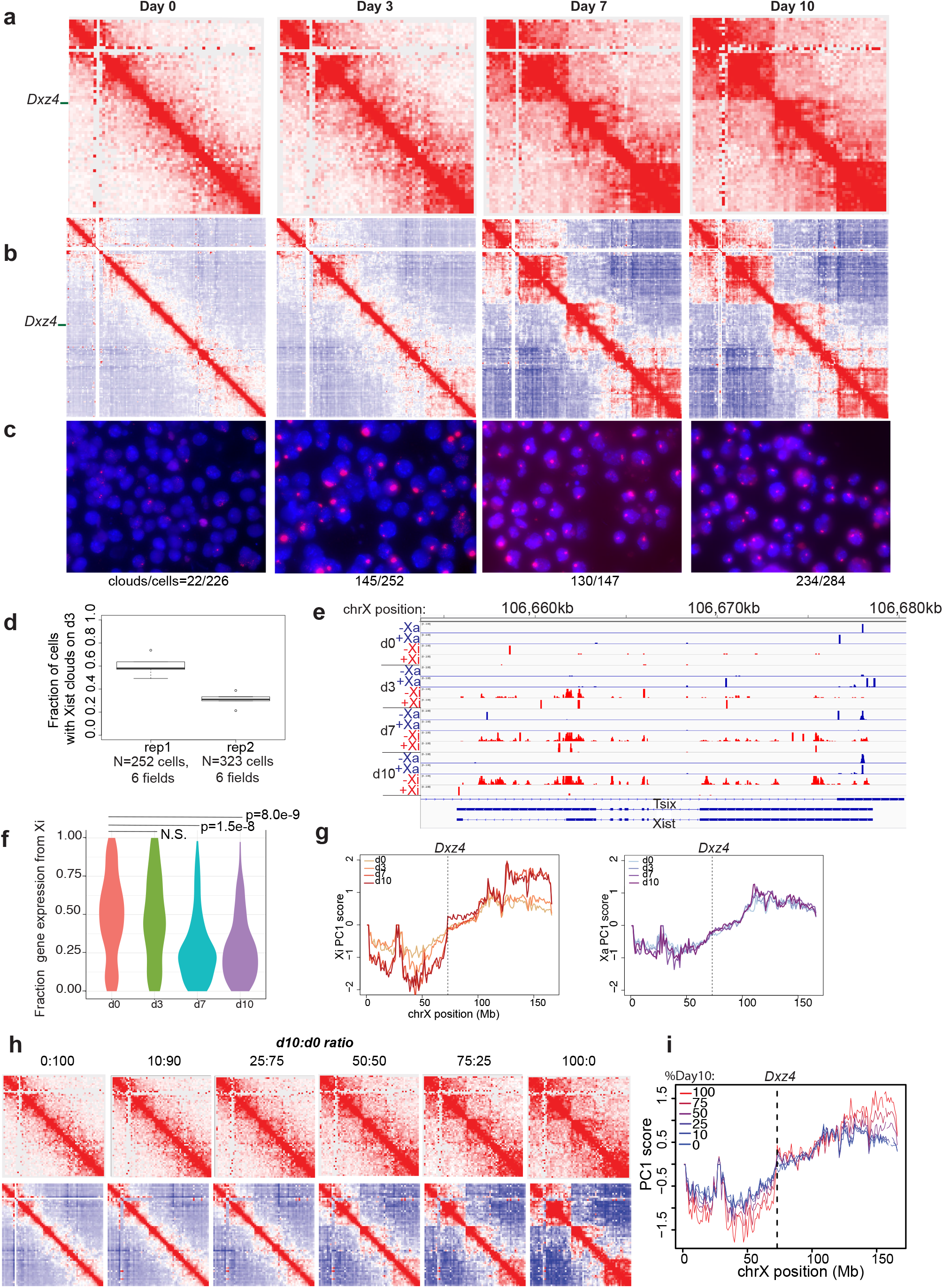
Dynamics of megadomain formation during XCI. (**a**) KR-normalized Hi-C matrices on future Xi (mus) in female ES cells on days 0, 3, 7 and 10 of differentiation (2.5 Mb resolution) (**b**) Pearson correlation of Hi-C matrices on days 0, 3, 7, 10 of differentiation (1 Mb resolution). (**c**) Xist RNA FISH on day 0, 3, 7, 10 of differentiation in *Tsix^TST/+^* ES cell line. **(d)** Fraction of cells showing Xist clouds on day 3 of differentiation in two biological replicates. Error bars represent standard deviation of counts over 6 fields in one imaging session. **(e)** Allele-specific expression from the Xi (red) or Xa (blue) over *Xist* in *Tsix^TST/+^* during differentiation. (**f**) Density plots of the number of X-linked genes with a given level of allelic expression from the Xi on day 0, 3, 7, 10 of differentiation in *Tsix^TST/+^*. Wilcox p-values comparing the mean allelic expression levels from the Xi between d0 and all other timepoints are indicated. (**g**) 1^st^ principal component of the 1Mb correlation matrix plotted for all bins on the future Xi (mus, left) and future Xa (cas, right) for Days 0, 3, 7, 10 of differentiation. Dotted lines correspond to the bin containing *Dxz4*. All Hi-C data in this figure are generated from merging together reads from two biological replicates. (**h**) In silico mixing experiment to determine the sensitivity of the Hi-C assay for detecting megadomains. KR-normalized Hi-C matrices at 2.5 Mb resolution (top) and Pearson correlation of Hi-C matrices at 1 Mb (bottom) for the Xi from data sets generated by mixing an indicated ratio of day 0 (d0, megadomain-negative) and day 10 (d10, megadomain-positive) datasets. (**i**) 1^st^ principal component of the 1Mb correlation matrix plotted for all bins on the Xi for datasets within varying ratios of d10:d0 reads.

On the other hand, analysis of day 7 cells revealed Xist expression in >80% of cells (Fig. 1c) and robust Xi silencing (Fig. 1f). It was at this timepoint that strong megadomains were first observed (Fig. 1a,b). Analysis of day 10 cells showed similarly strong Xist expression, Xi silencing, and megadomain formation. To quantify megadomain signals, we computed the Pearson correlation for the Xi contact maps and performed principal component analysis (PCA). On day 7 and day 10, there was a sharp transition in the 1^st^ principal component score (PC1) at *Dxz4*, indicating changed interaction patterns on each side of *Dxz4* at later timepoints but not in days 0 or 3 (Fig. 1g), consistent with appearance of megadomains. By contrast, the PC1 score distribution for the Xa was nearly identical for all timepoints without a sharp transition at *Dxz4* (Fig.1g). In addition, the sharp transition in the PC1 curve is a valid measure of megadomain strength, as our mixing experiment showed that the slope of the curve at *Dxz4* was directly proportional to the fraction of day 10 cells in the mixing experiment (Fig S2e). The dynamics of megadomain formation were highly reproducible between two biological replicates (Fig. S3a). Taken together, these data suggest that megadomains do not precede XCI and appear either concurrently with or (more likely) only after Xist has spread and silenced the Xi.

### Time course of TAD attenuation on the Xi

Recent analyses indicate that TADs are not abolished on the Xi but are instead attenuated^24,29^. Here we investigate the time course of TAD attenuation during XCI. To enrich for interactions and obtain higher resolution allele-specific contact maps, we performed Hi-C^2^, a variation of the Hi-C protocol that focuses analysis on defined regions through hybrid capture using high density probe sets^6^. We investigated ~1.5 Mb regions around (i) *Dxz4* to assess the behavior of the strong megadomain border and (ii) the TAD harboring the disease locus and inactivated gene, *Mecp2*, to examine how topological domains are weakened during XCI.

Interestingly, despite megadomains appearing only late during the XCI time course, the *Dxz4* region showed a strong boundary during all time points and on both alleles (Fig. 2a,b). Therefore, *Dxz4* acts as a border irrespective of XCI status and presence/absence of megadomains. At days 7 and 10, the proximal TAD flanking *Dxz4* strengthened and expanded on the Xi but not the Xa (Fig. 2a,b), leaving only two “boxes” on either side of *Dxz4* by day 10, rather than the patchwork of smaller sub-TADs present on the Xa and Xi at earlier timepoints. This finding indicated that the emergence of a megadomain correlates with increased insulation by *Dxz4*, indicating that *Dxz4* insulates interactions at increasingly larger distances when megadomains form. The temporal dynamics of chromatin conformation surrounding *Dxz4* were quite similar in two independent replicates of the differentiation timecourse and Hi-C^2 enrichment (Fig. S3b).

**Figure 2:**
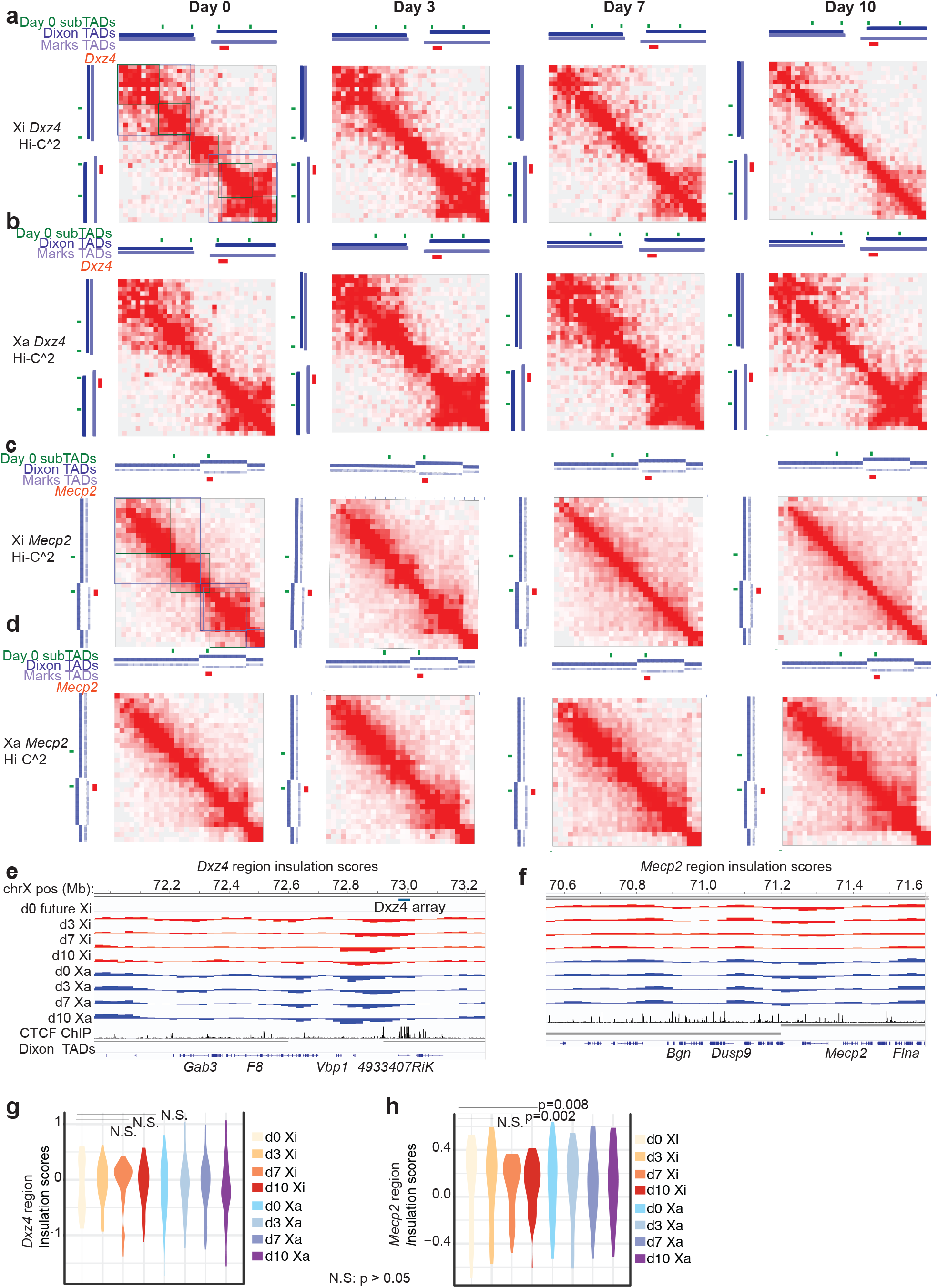
Dynamics of TADs and sub-TADs during XCI in the *Mecp2* and *Dxz4* regions. (**a,b**) Hi-C^2^ contact maps around *Dxz4* (mm9 coordinates chrX:71,832,976-73,511,687) and on the future Xi (**a**) or Xa (**b**) on days 0, 3, 7, 10 of ES differentiation (50 kb resolution). (**c,d**) Hi-C^2^ contact maps around *Mecp2* (mm9 coordinates chrX:70,370,161-71,832,975) and on the future Xi (**c**) or Xa (**d**) on Days 0, 3, 7, 10 of differentiation (50 kb resolution). Green bars indicate positions of sub-TAD borders determined from 25 kb d0 comp Hi-C^2^ matrices; dark blue track shows Dixon et al^2^. TAD calls in mESCs, light blue track shows Marks et al^56^. TAD calls in mESCs, red bars indicate positions of either *Dxz4* or *Mecp2*. In addition, the regions of the contacts corresponding to sub-TADs (green), Dixon et al. TADs (dark blue), Marks et al. TADs (light blue) have been indicated with boxes on the day 0 Xi contact maps for reference. (**e,f**) Insulation scores across the *Dxz4* region (**e**) or Insulation scores across the *Mecp2* region (**f**). Insulation scores on the Xa are in blue and insulation scores on the Xi are red. For reference, CTCF ChIP-seq in d0 F1-2.1 mESCs (black) and TAD calls from Dixon et al. (grey) are shown^2^. (**g,h**) Violin plots showing the distributions of insulation scores across the *Dxz4* region (**g**) and *Mecp2* region (**h**). All data in this figure are generated from merging together reads from two biological replicates. Note: to generate violin plots and evaluate the significance of differences in variance between timepoints we excluded the 6 bins on each edge of the Hi-C^2^ region because the regions needed to calculate insulation score fall partly outside the Hi-C^2^ region and have far lower read counts than sequences targeted by the capture probes.

Within the region containing *Mecp2*, Hi-C^2 contact maps showed that X^mus^ and X^cas^ behaved similarly on days 0 and 3, in that both were organized into several sub-TADs (Fig. 2c,d). However, once Xist spread over X^mus^ and the Xi formed as a consequence on days 7 and 10, both TAD and sub-TAD organization become obscured compared to the Xa, where these domains stayed similar to earlier timepoints. In contrast to a previous analysis performed in neural progenitor cells (NPCs)^22^, we did not observe the persistence of a small domain around *Mecp2* in differentiating female ES cells. The loss of domain organization in the *Mecp2* region was observed in two distinct biological replicates (Fig. S3c).

To quantify these changes, we computed insulation scores using standard methods ^22,41^ (Fig. 2e,f). In brief, insulation scores quantify how strongly a given locus acts as a border^41-43^, and are calculated by running sliding windows across a chromosomal region and measuring the log ratio of reads crossing over a locus to reads neighboring a locus (Fig. S3d). Loci in the interiors of a TAD would be expected to have similar numbers of cross-over interactions and local interactions on each side, leading to insulation scores near zero, whereas loci at borders would register as a local minimum of crossover interactions than local interactions, leading to strong negative insulation scores at domain boundaries. We observed several interesting facets of the insulation score curves on the Xa and Xi at *Dxz4* and *Mecp2* during the differentiation timecourse. There was a strong decrease in insulation scores near *Dxz4* on both alleles at all timepoints, consistent with *Dxz4* acting as a boundary throughout differentiation (Fig. 2e,g). The variance of the insulation scores is a measure of the global strength of insulation, with smaller variance corresponding to weaker insulation^44^. Near *Dxz4*, the variance was slightly smaller on the Xi than the Xa for all timepoints. There was no statistically significant difference in variance of insulation scores on the Xi on day 0 compared with the variance insulation scores on the Xi for the later timepoints (pairwise F-test p-values >0.05 for all comparisons between day 0 Xi and later Xi timepoints) and this observation held across two biological replicates (Fig. 2g, Fig. S3e). Thus, the *Dxz4* region is a strong boundary regardless of XCI status, but that *Dxz4* insulates interactions from increasingly larger distances to form megadomains.

The *Mecp2* region showed a different pattern of insulation score changes across differentiation. The distribution of insulation scores across the *Mecp2* region significantly narrowed on days 7 and 10 on the Xi but not Xa (Fig. 2h). Indeed, across the *Mecp2* region, the variance on the day 7 or day 10 Xi was significantly lower than on day 0 (day 7 vs day 0 p-value=0.008443; day 10 vs day 0 p-value=0.001819). There was no significant difference in the variance on the Xi between day 0 and day 3 (p=0.7369). This clear decrease in the variance on the Xi at day 7 and day 10 relative to d0 was observed in two biological replicates (Fig. S3f). These results indicate that TAD and sub-TAD structures of the *Mecp2-*containing TAD region are reduced in strength in the same timeframe that megadomains are gained.

### *Dxz4* is necessary but not sufficient for megadomain formation

There presently exist three deletions containing *Dxz4/DXZ4* — two in mouse^22,25,29^, one in human^25^. One of the mouse deletions^22^ is a large deletion that contains more than just the noncoding element, *Dxz4/DXZ4* (Fig. 3a). We generated a new deletion of *Dxz4* and its flanking sequences that left untouched a small cluster of CTCF motifs with very high CTCF coverage and an unusual satellite repeat (Fig. 3a and Fig. S4a)^28^, both of which were deleted in a previous 200-kb *Dxz4* deletion. We generated a smaller 100-kb deletion spanning *Dxz4* (*Dxz4^∆100^*) and validated our deletion by Sanger sequencing, by DNA fluorescence in situ hybridization (FISH) with a probe internal to the deleted region, and by genomic DNA sequencing to examine read distributions over the deleted region (Fig. S4, Methods). Importantly, to distinguish Xi from Xa, we performed the deletion analysis in *Tsix^TST^*/+ mESCs.

**Figure 3:**
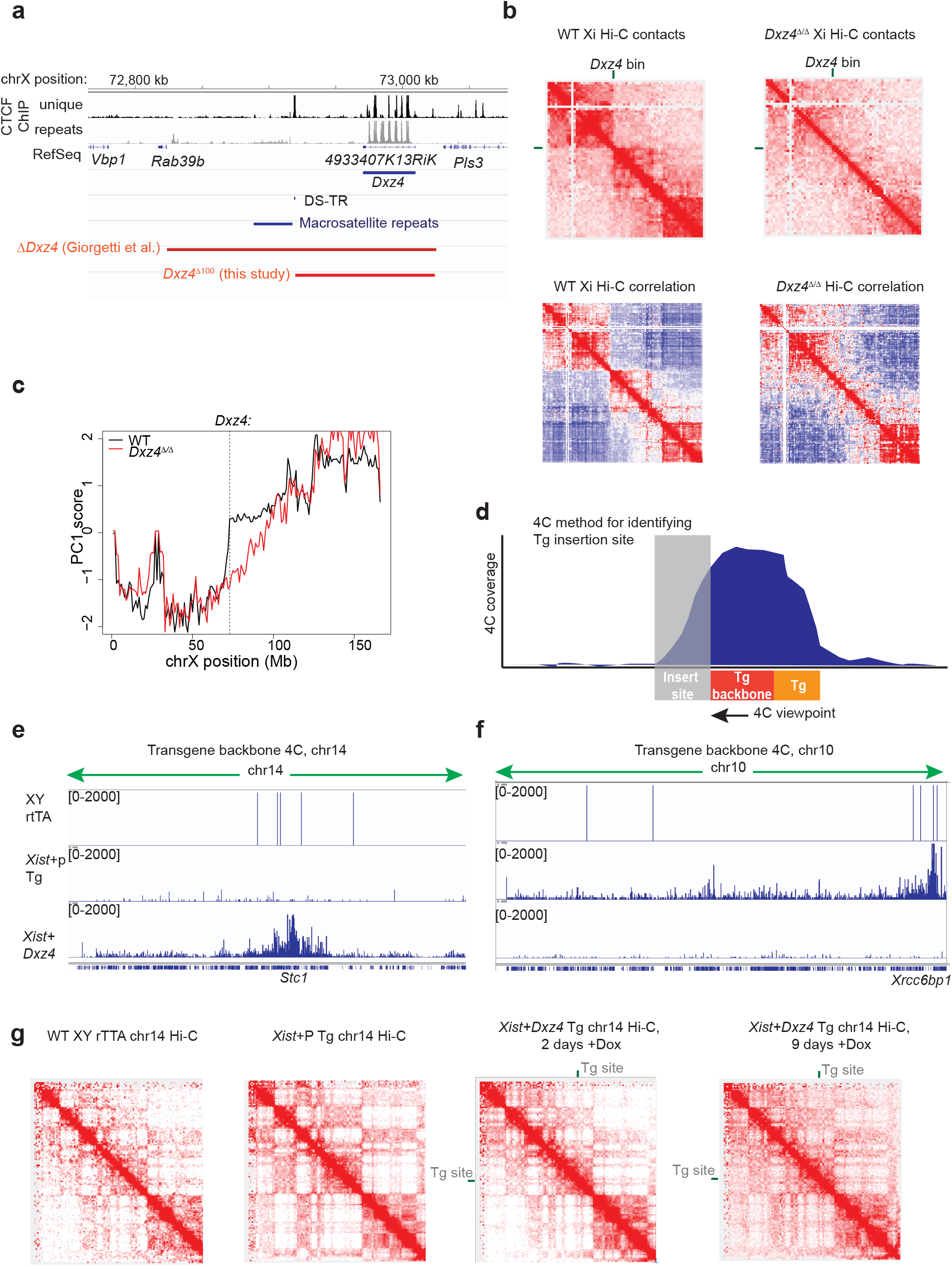
*Dxz4* is necessary but not sufficient for megadomain formation. (**a**) Schematic of prominent features in the *Dxz4* region and deletions generated in this study and previous studies. (**b**) Top: Contact maps for the Xi at 2.5Mb resolution for wild-type (left) and *Dxz4*^∆/∆^ (right) cells on d10 of differentiation. Bottom: Pearson correlation matrices at 1Mb resolution for the wild-type (left) and *Dxz4*^∆/∆^ Xi (right). Heatmaps generated by merging reads together from two biological reploicates. (**c**) 1^st^ principal component of the 1Mb correlation matrix plotted for wild-type (black) and *Dxz4*^∆/∆^ (red) across all bins on the Xi. The dotted line corresponds to the bin containing *Dxz4*. (**d**) Strategy for using 4C to localize the transgene insertion site. (**e,f**) 4C interaction profiles using a viewpoint in the backbone of an *Xist* transgene in either wild-type (top), *Xist* only Tg (middle) or *Xist+Dxz4* Tg (bottom) lines across chr14 (**e**) or chr10 (**f**). (**g**) Hi-C contact maps for chr14 at 1Mb resolution in either wild-type (left), Xist only Tg (middle left), *Xist+Dxz4* Tg Dox induced for 2 days (middle right) or 9 days (right). The position of the transgene insertion site at ~69.8Mb (near *Stc1*) is indicated by a green bar in the *Xist+Dxz4* Tg contact maps.

To test the impact of removing *Dxz4*, we differentiated wild-type and homozygously deleted (*Dxz4^∆/∆^*) cells for 10 days and performed Hi-C. Whereas wild-type cells showed strong megadomains, *Dxz4^∆/∆^* cells showed disrupted megadomain structures in Hi-C contact maps (Fig. 3b) and corresponding Pearson correlation maps. Most prominently, the sharp border around *Dxz4* was eliminated, though some intra-megadomain interactions remained on either side of the deletion. A disrupted megadomain border was confirmed by loss of the sharp transition in PC1 score around the *Dxz4* locus (Fig. 4c). This effect was observed in two biological replicates (Fig. S4d). These results are in agreement with prior reports^22,25,29^ that the 200-300 kb region around *Dxz4* is required for megadomain organization. Additionally, our work delineates the required region to a 100-kb domain containing the *Dxz4* tandem repeat itself (as opposed to the CTCF motif cluster and the proximal satellite repeats).

**Figure 4:**
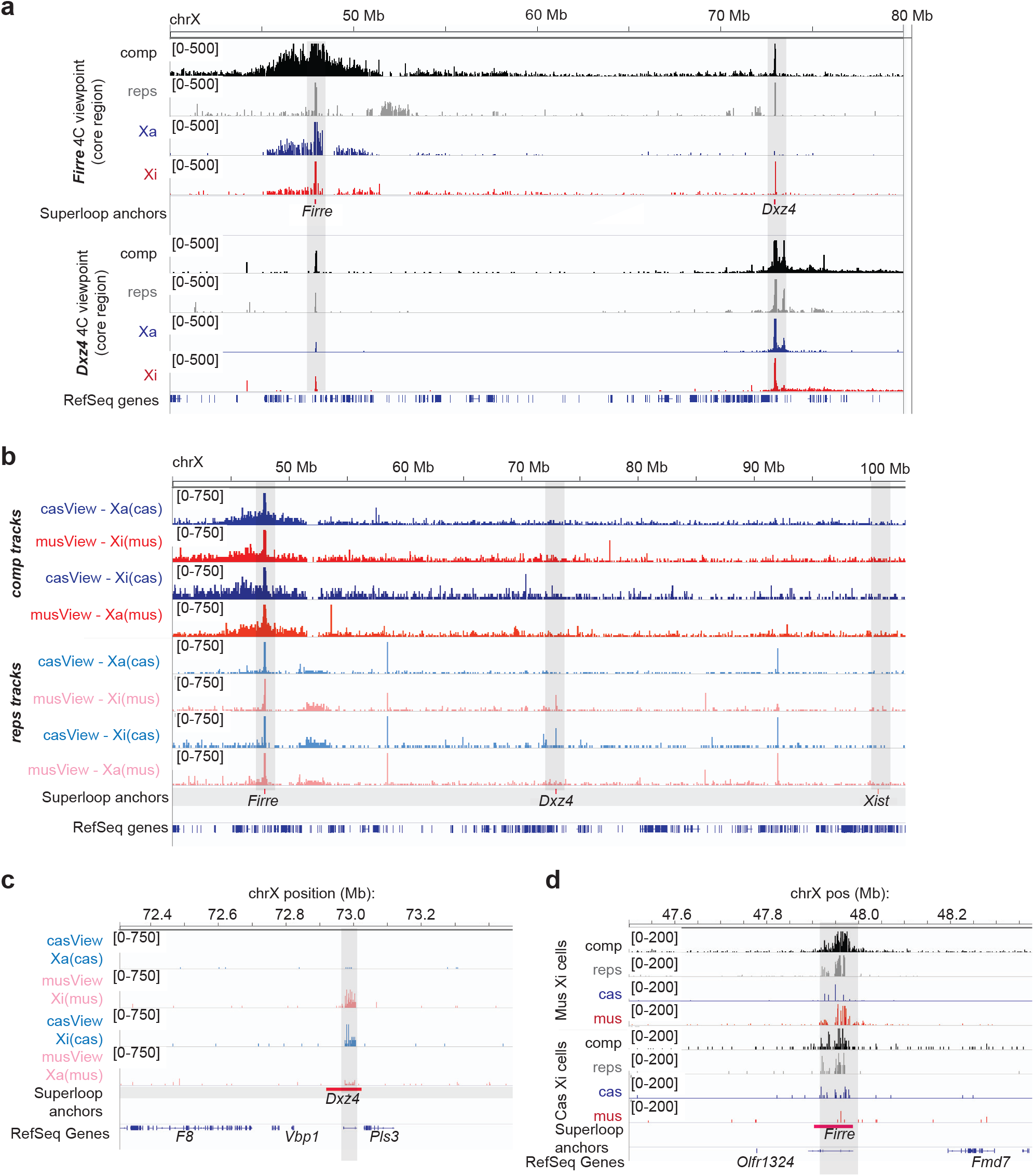
*Dxz4, Firre*, and Xi-specific superloops. (**a**) Reciprocal interaction between *Dxz4* and *Firre* in post-XCI fibroblasts. Top: 4C interaction profiles with the core of the *Firre* tandem repeat as the viewpoint. Bottom: 4C interaction profiles with the core of the *Dxz4* tandem repeat as the viewpoint. Black: all unique reads (comp). Grey: repetitive (non-unique) reads (comp). Blue: Xa (cas) specific reads. red: Xi (mus) specific reads. (**b**) 4C from a unique, allele-specific viewpoint at the 3’ end of *Firre* in cells with either an inactive mus (cas Xa/mus Xi) or an inactive cas (cas Xi/mus Xa). Red: interaction profile for unique (comp) reads on the mus allele. Blue: interaction profile for unique (comp) reads on the cas allele. Pink: interaction profile for non-unique (reps) reads from the mus allele. Light Blue: interaction profile for non-unique (reps) reads from the cas allele. Positions of *Firre*, *Dxz4* and *Xist* are shaded light grey and indicated with red bars. (**c**) 4C interaction profile over *Dxz4* from a unique, allele-specific viewpoint at the 3’ end of *Firre* in cells with either a mus Xi (top tracks) or a cas Xi (bottom tracks). Light blue: interaction profile from the cas *Firre* viewpoint. Pink: interaction profile from the mus *Firre* viewpoint. (**d**) 4C interaction profile over *Firre* using the *Dxz4* core viewpoint in cells either with a mus Xi (top tracks) or a cas Xi (bottom tracks). Black: all unique (comp) reads. Grey: non-unqiue (repetitive) reads. Blue: cas reads. Red: mus reads. Note: **(c)** and **(d)** zoom into *Dxz4* and *Firre* respectively to better highlight the allele-specific superloop interactions.

Given its necessity for megadomain organization, we asked whether *Dxz4* is also sufficient to form a megadomain on an autosome when Xist RNA is expressed in *cis*. We co-transfected a dox-inducible full-length *Xist* construct along with a BAC containing *Dxz4* into male fibroblasts and used RNA and DNA FISH to identify *Xist*-inducible clones (Fig. S5a) where both *Xist* and *Dxz4* had co-inserted (XPDxz4.4; Fig. S5b). To localize the transgene, we adapted the 4C technique^45,46^ that is ordinarily used to view 3D interactions from a single locus. We reasoned that, by placing the 4C viewpoint anchor at the transgene backbone, we could map the transgene through the pattern of *cis-*interactions on the same chromosome (Fig. 3d), as interaction frequencies are typically highest near the viewpoint position. This fact has previously been used to aid genome assembly^47-49^. Indeed, in addition to interaction peaks at *Xist* and *Dxz4* as expected (Fig. S5c), the only other strong 4C peak in the genome appeared on chr14 near *Stc1* (Fig. 3e). This finding contrasts with both a control *Xist-*only transgene line which showed a peak only on chr10 (Fig. 3f) and the parental rtTA fibroblasts which showed no peaks anywhere in the genome. We confirmed insertion of both *Dxz4* and the *Xist* construct into Stc1 by observing co-localization between *Stc1*, *Xist* and *Dxz4* DNA FISH probes at one spot in the transgenic cell line (Fig. S5d).

To test whether induction of Xist expression could induce megadomain formation at *Dxz4* ectopically, we induced *Xist* and performed Hi-C to determine whether co-insertion of *Xist* and *Dxz4* induced formation of megadomains on transgenic chr14 in fibroblasts. We induced Xist for 2 days because a previous report suggested that induction of Xist from the male X for 2 days was sufficient to at least initiate megadomain formation^22^. No megadomains formed and the overall chr14 contact maps looked highly similar to the non-transgenic and *Xist*-only controls (Fig. 3g). We then extended the time frame and induced for 9 days, given that our ES cell time course suggested that several days of Xist upregulation were needed to form megadomains on the Xi. Still, no megadomains formed in these post-XCI cells (Fig. 3g). To assess whether *Xist* and *Dxz4* could do so in cells undergoing de novo XCI, we attempted three times to create the *Xist-Dxz4* transgene line in a female ES cell background, but such a line could not be generated, due to potential lethal consequences of the *Xist* transgene. These results indicate that *Xist* and *Dxz4* together are not sufficient for megadomain formation in a cell line that had already undergone XCI (fibroblasts). We conclude that *Dxz4* and *Xist* expression are necessary but not sufficient for megadomain formation in post-XCI cells.

### *Dxz4, Firre*, and Xi-specific superloops

In addition to serving as the border between the megadomains, *Dxz4* has been shown to form extremely long (>10 Mb) looping interactions with other loci on the human Xi^1,25,27^. To further dissect the role of *Dxz4* in establishing the large-scale structure of the Xi, we performed 4C using a viewpoint within the core of the *Dxz4* tandem repeats in post-XCI fibroblasts to identify interacting loci that may be important for helping to establish the unique structure of the mouse Xi. *Dxz4* generally interacted with the chromosome telomeric to *Dxz4* and formed few long-range interactions towards the centromeric side of the chromosome (Fig. 4a, Fig. S6a). However, *Dxz4* interacted strongly with another non-coding tandem repeat, *Firre* (Fig. 4a). The two loci formed an extremely strong loop despite the fact that *Firre* is 25 Mb centromeric to *Dxz4*. The strength of their interaction was equivalent to that of two loci separated by < 200 kb (data not shown). To verify the *Dxz4:Firre* interaction, we performed a reciprocal 4C using a viewpoint within the core of the *Firre* tandem repeats and confirmed a strong reciprocal interaction (Fig. 4a). On the Xa, *Firre* also formed a broad domain of interactions with nearly all sequences within several Mb of itself, as reported for the Firre RNA contact map previously^30^. In contrast to this prior study, however, we did not observe any evident interchromosomal contacts from either Xa or Xi allele in fibroblasts.

Our allele-specific analysis revealed that the *Dxz4:Firre* interaction is primarily detected on the Xi (mus) allele. Indeed, when we repeated this reciprocal 4C experiment in another hybrid fibroblast line that chose to inactive X^cas^, the *Dxz4:Firre* interaction was detected on X^cas^, rather than X^mus^. For these experiments, we used both a unique 4C anchor in the 3’ flanking region of *Firre* that provides allelic information, as well as an allele-agnostic anchor in the core of *Firre* repeat. Because *Dxz4* and *Firre* are both highly repetitive, we also examined multiply-aligning reads. The *Firre-Dxz4* interaction was only observed on the Xi regardless of whether the Xi was the mus or cas chromosome (Fig. 4b-d). Thus, *Firre* and *Dxz4* formed an Xi-specific superloop conserved between mouse and primate^1,25^.

Other superloops have been identified on the human Xi using high-resolution Hi-C. *FIRRE, DXZ4, XIST, ICCE* and *X75* are all repetitive loci that bind CTCF on the Xi in human cells, and all form long-range interactions with each other^1,25^. We examined whether these superloops also occur in mouse cells. Indeed, in addition to *Firre*, we observed elevated interaction frequencies between *Dxz4* and a region spanning *Xist* to *Ftx* (Fig. S6a,b) and a region syntenic with human *X75* (Fig. S6a,c). However, the *Xist-Dxz4* and *x75-Dxz4* contacts were less prominent than the *Firre-Dxz4* contact.

### *Firre* is predominantly expressed from the Xa

Previous reports have suggested that *Firre* escapes from X-inactivation^30,32,50^, and that Firre RNA is necessary for *Xist* localization and deposition of H3K27me3 on the Xi^32^. By allele-specific RNA-seq, we observed Firre reads from both Xa and Xi during differentiation, and expression appeared to be predominantly though not exclusively exonic Fig. 5a,b). Because *Firre* is highly repetitive, SNP calls may be not be fully reliable. To confirm allele-specific expression, we used genetic means to examine expression from the Xa and Xi. With allele-specific guide RNAs, we generated an Xi-specific *Firre* deletion in *Tsix^TST^/+* (“*Firre*^*Xi*∆/+^ clone D1”), an Xa-specific Firre deletion in *Tsix^TST^/+* (“*Firre*^Xa∆/+^ clone H6”), and a homozygous deletion (“*Firre*^∆/∆^”) in female ES cells (Fig. S7). We then measured *Firre* expression on day 10 of differentiation using 4 published sets of primers^30,32^ and one new intronic primer set and deduced the expressed allele(s) by examining differences in expression pattern between the reciprocal heterozygous clones. First, by quantitative RT-PCR of wild-type female ES cells, we inferred that Firre expression was expressed at <10% of Xist RNA overall. The variability between amplicons suggested that there could be multiple isoforms of Firre (Fig. 5d). Second, deleting *Firre* on the Xi abolished expression of the two lowest expressed amplicons (Fig. 5e,f). By contrast, deleting *Firre* on the Xa abolished expression of the two most highly expressed exonic amplicons and the intronic amplicon. Finally, a homozygous deletion abolished all expression measured from these primer pairs.

**Figure 5:**
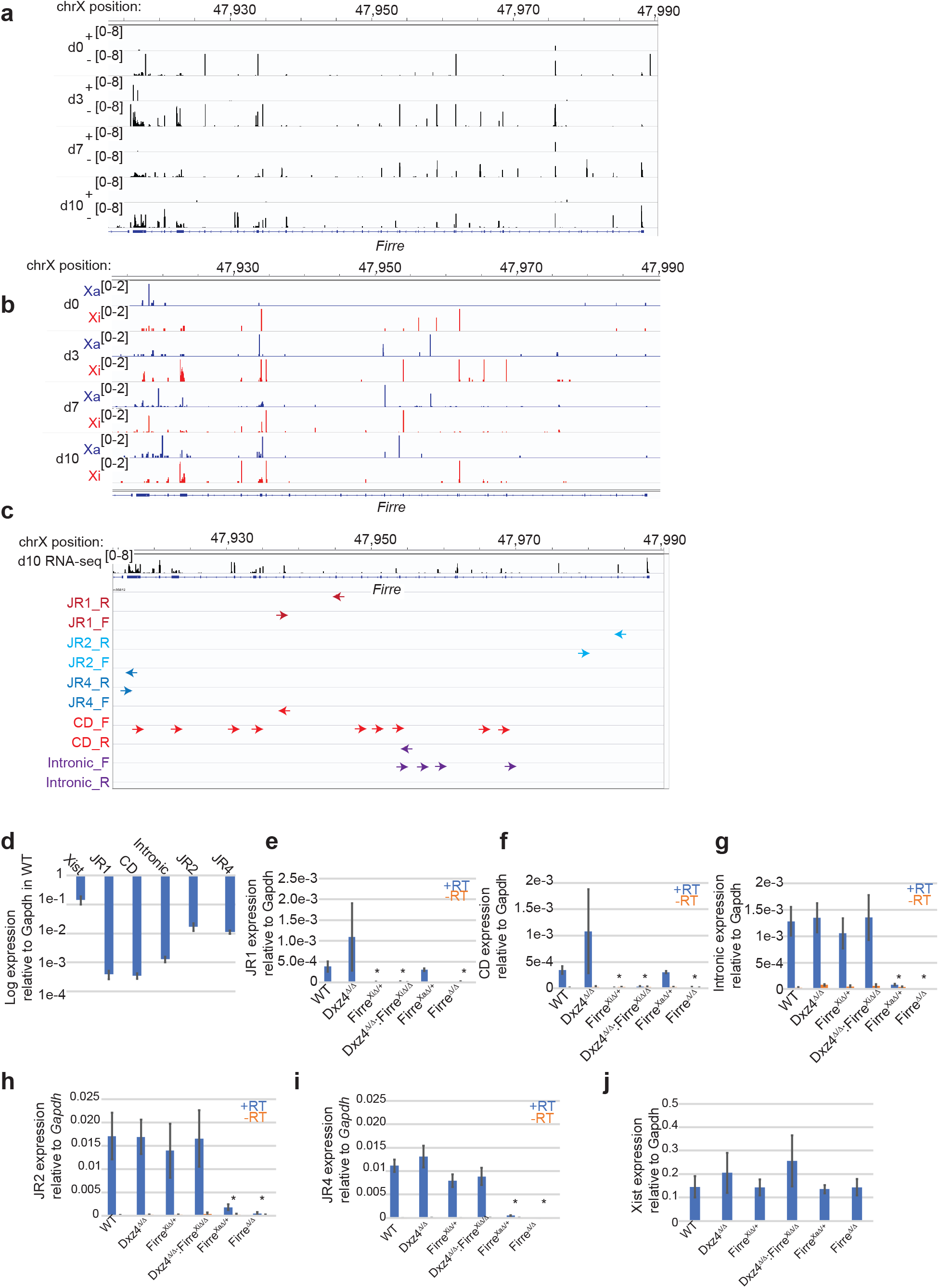
*Firre* is predominantly expressed from the Xa in differentiating female ES cells. (**a**) Expression over *Firre* during differentiation. (**b**) Allelic expression over *Firre* during differentiation, red=Xi reads, blue=Xa reads. (**c**) Positions of Firre primers within the *Firre* locus. (**d**) Expression of Xist or Firre amplicons in d10 wild-type cells normalized to Gapdh (logarithmic scale). (**e**-j) Expression normalized to Gapdh measured with JR1 (**e**), CD (**f**), intronic (**g**), JR2 (**h**), JR4 (**i**) Firre primers or Xist primers (**j**) in WT, *Dxz4*^∆/∆^ *Firre*^Xi∆/+^, *Dxz4*^∆/∆^:*Firre*^Xi∆/+^, *Firre*^Xa∆/+^ and *Firre*^∆/∆^ cells. Blue bars are +RT, orange bars are -RT. Asterisks indicate a statistically significant (p < 0.05, t-test) difference between normalized expression levels in WT and normalized expression levels in the deletion. Error bars show standard error of the mean, 3 biological replicates.

To determine if *Dxz4* influences *Firre* expression, we also performed RT-PCR in the *Dxz4*^∆/∆^ cell line. No changes were evident, indicating that the *Dxz4*-*Firre* superloop does not impact transcriptional regulation of *Firre* (Fig. 5e-i). Finally, to examine whether either repeat locus regulates *Xist* expression, we performed RT-PCR in cell lines carrying deletions of either *Firre* or *Dxz4*, and also generated and tested a double ES cell knockout of *Dxz4* and *Firre* on the Xi carrying the *Tsix^TST^* allele (*Dxz4*^∆/∆^:*Firre*^*Xi*∆/+^)(Fig. S7, Methods). None of these deletions affected *Xist* expression (Fig. 5j). Together, our results suggest that different isoforms of Firre RNA are expressed from the Xa and Xi, but expression may predominate on the Xa. The results also indicate that the superloops are irrelevant for expression of *Dxz4*, *Firre*, and *Xist*.

### *Firre* is not strictly required for megadomain formation, but works together with *Dxz4* to strengthen megadomains

Prior work had not examined whether Xi superloops contribute to the formation of the megadomain boundary at *Dxz4*. Our *Firre* deletion lines allowed us to use Hi-C to test whether loss of the other anchor in the *Dxz4-Firre* superloop perturbed megadomains. To exclude possible *trans-*effects relating to *Firre* RNA^51^ expressed from the Xa and to focus on the role of *Firre* locus as a superloop anchor, we conducted all analysis in our Xi-specific *Firre* deletion. We performed Hi-C on day 10 of differentiation in wild-type, *Firre*^Xi∆/+^ and *Dxz4*^∆/∆^:*Firre*^Xi∆/+^ cells. Importantly, despite disruption to the other side of the *Dxz4-Firre* superloop, the *Firre*^Xi∆/+^ cells retained the sharp megadomain boundary on the Xi. However, in two biological replicates, the intra-megadomain interactions appeared attenuated compared to wild-type (Fig. 6a,b; S8). Plotting PC1 scores for all bins on the Xi showed a sharp transition in PC1 score at *Dxz4* in wild-type and *Firre*^Xi∆/+^ (Fig. 6c), in agreement with the continued presence of a bipartitle structure. Thus, *Firre* may influence the strength of interactions within each megadomain, but is not strictly required for formation of the *Dxz4* border and the bipartite mega-structures. Given the phenotypes of the *Dxz4* and *Firre* single deletions, we predicted that the double deletion would obliterate all trace of megadomains. Indeed, the *Dxz4*^∆/∆^:Firre^Xi∆/+^ cells showed a loss of the sharp border at *Dxz4* and a depletion of intra-megadomain interactions (Fig. 6a-c). Based on the Pearson correlation heatmap and PC1 analysis, the double deletion may potentially reduce the intra-megadomain interactions on either side of *Dxz4* more than in the *Dxz4^∆/∆^* single deletion, though the sharp border at *Dxz4* was lost in both cases (Fig. 3,S4 versus Fig. 6, S8). We conclude that *Firre* is not absolutely required for megadomain formation. However, *Firre* and the *Firre-Dxz4* superloop may work together with *Dxz4* to strengthen the megadomain structure.

**Figure 6:**
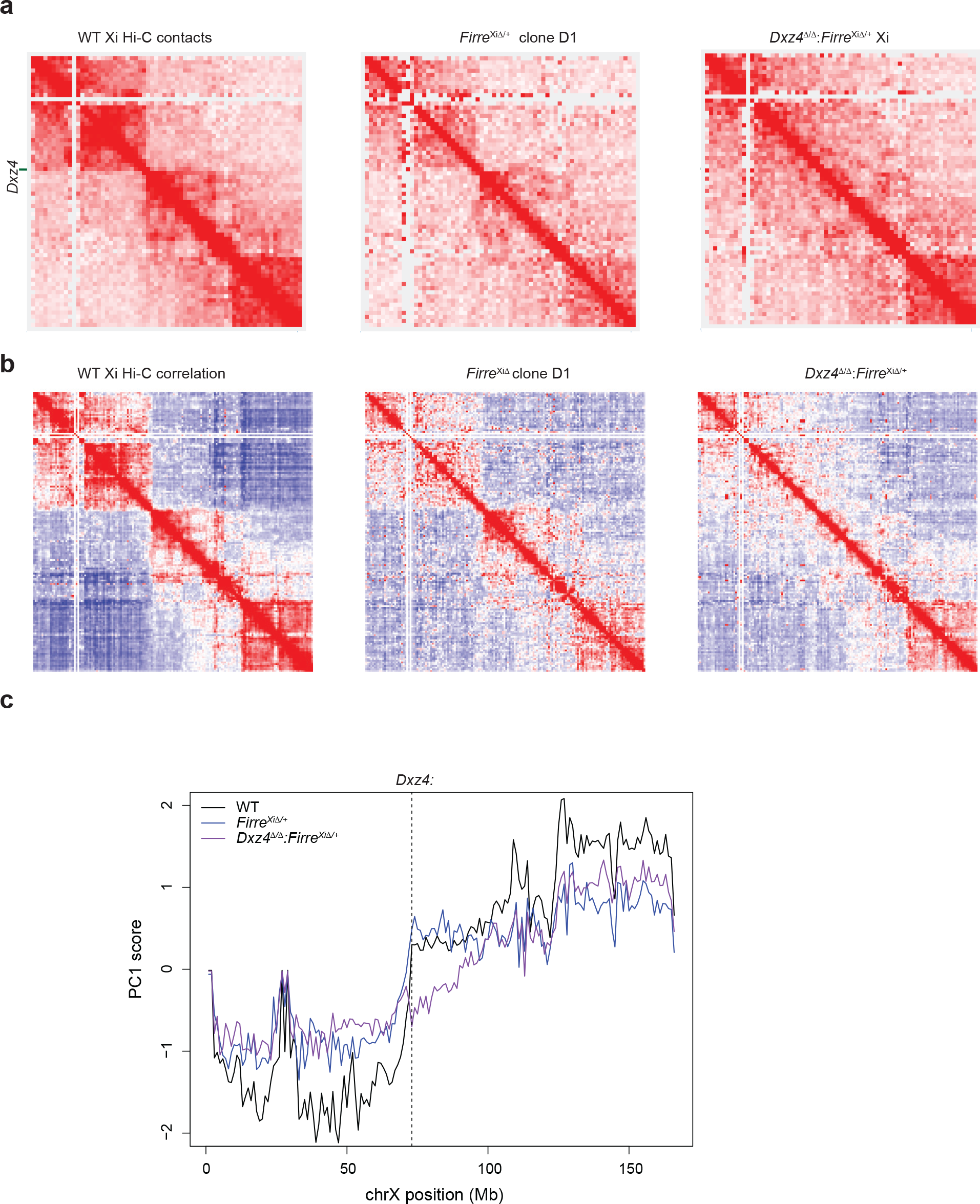
*Firre* is not required for megadomain formation but deleting *Firre* weakens megadomains. (**a**) Contact maps for the Xi at 2.5Mb resolution for wild-type (left), *Firre*^Xi∆/+^ (center) and *Dxz4*^∆/∆^:*Firre*^Xi∆/+^ cells on d10 of differentiation. (**b**) Pearson correlation matrices at 1Mb resolution for the wild-type (left) *Firre*^Xi∆/+^ (center) and *Dxz4*^∆/∆^:*Firre*^Xi∆/+^. (**c**) 1^st^ principal component of the 1Mb correlation matrix plotted for wild-type (black) and *Firre*^Xi∆/+^ (blue) and *Dxz4*^∆/∆^:*Firre*^Xi∆/+^ (purple) across all bins on the Xi. The dotted line corresponds to the bin containing *Dxz4*. All data in this figure are generated from merging together reads from two biological replicates.

### Megadomains and superloops can be uncoupled from XCI and escape

Recent studies have diverged on the effect of deleting the *Dxz4* region on XCI, as one study suggested a loss of escape from XCI for many escapees in mouse NPCs^22^ and another suggested a partial loss of H3K27me3 on the human fibroblast Xi^25^. A prior report also suggested that knockdown of Firre RNA disrupts localization of the Xi to the nucleolus and maintenance of H3K27me3 on the Xi^32^. Here we assessed the effect of deleting *Dxz4*, *Firre*, or both on various aspects of XCI. First, we examined effects on the Xist RNA cloud that normally forms over the Xi, but observed no obvious changes in Xist cloud morphology or number of cells exhibiting *Xist* upregulation on day 10 of differentiation in *Dxz4^∆/∆^*, *Firre*^Xi∆/+^, *Firre*^Xa∆/+^, *Dxz4^∆/∆^*:*Firre*^Xi∆/+^ or *Firre*^∆/∆^ (Fig. 7a,b, Fig. S9a,b) versus wildtype female cells. There was also no effect on the characteristic localization of Xist RNA/Xi to the perinucleolar region^52^(Fig.7a,b, Fig. S9a,b). We also observed no difference in the enrichment of the H3K27me3 repressive mark on the Xi after 7 days of differentiation in any of the *Dxz4* or *Firre* deletions (Fig. 7a,c) or after 10 days in the *Dxz4* and Xi-specific *Firre* deletions (Fig. S9a,c). This suggests that neither *Firre* nor *Dxz4* is required for Xist to be expressed, localized, and deposit H3K27me3 on the Xi. To test whether there is a partial loss of H3K27me3 across a macroscopic region of the Xi, as observed in a human DXZ4 deletion^25^, we produced metaphase spreads in WT and *Dxz4*^∆/∆^ cells and performed H3K27ac and H3K27me3 immunofluoresence to visualize the Xi. The Xi stood out as the chromosome with almost no H3K27ac signal and very strong H3K27me3 signal^53^ (Fig. S9d). However, we observed no obvious difference between the H3K27me3 banding pattern on the WT or *Dxz4*^∆/∆^ Xi in metaphase spreads, suggesting no loss of H3K27me3 across a large region of the mouse Xi following *Dxz4* deletion (Fig. S9e).

**Figure 7:**
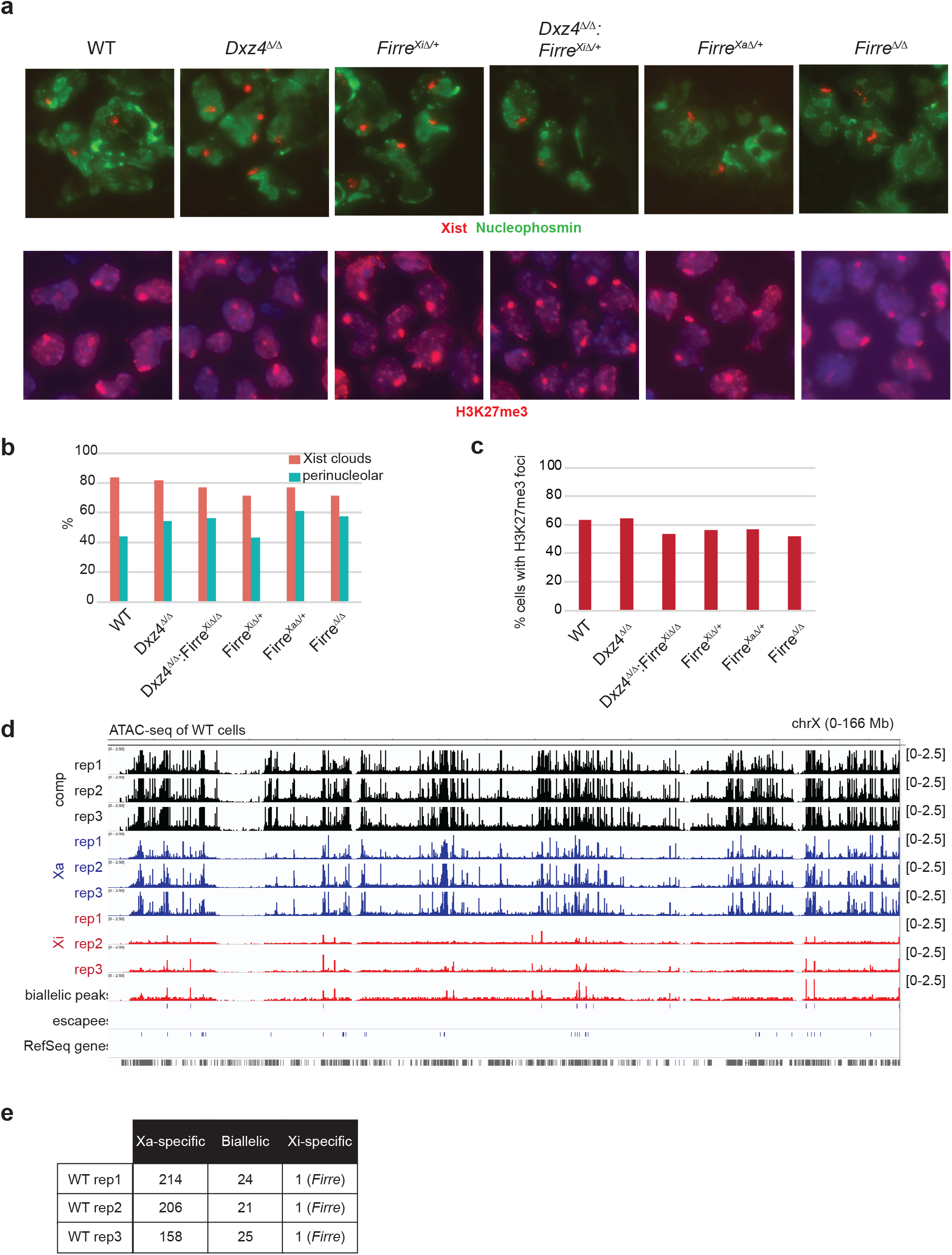
*Dxz4* and *Firre* do not affect Xi localization and accessiblity. (**a**) Xist RNA FISH (red) combined with nucleophosmin immunofluorescence (green) (top) and H3K27me3 IF (bottom) in WT, *Dxz4*^∆/∆^ *Firre*^Xi∆/+^, *Dxz4*^∆/∆^:*Firre*^Xi∆/+^, *Firre*^Xa∆/+^ and *Firre*^∆/∆^ cells. (**b**) Fraction of cells with Xist clouds (red) and fraction of Xist clouds in the perinucleolar space (cyan). (**c**) Fraction of cells with an H3K27me3 focus. (**d**) ATAC-seq coverage across the X (all unique reads, black), Xa (blue) and Xi (red) in WT cells, 3 biological replicates. (**e**) Allelic status of ATAC peaks in WT cells.

We next used ATAC-seq^54^ to assay chromatin accessibility on the Xi. In wild-type cells, ATAC signal was indeed heavily skewed towards the Xa, with >85% of all peaks on the X binding specifically to the Xa (Fig. 7d). There were ~20 biallelic sites (e.g., promoters of escapee genes) and only one Xi-specific peak (at *Firre*) (Fig. 7e). If *Dxz4* impairs X-chromosome accessibility as previously proposed for escapee genes^22^, we would expect nearby biallelic peaks near in wild-type to become Xa-specific in the deletion. On the other hand, if *Dxz4* or *Firre* were required to inhibit chromatin accessibility, we would expect many ATAC-peaks to appear on the Xi near genes subject to XCI. Significantly, the overall ATAC-seq patterns were highly similar between wild-type and all mutant genotypes — *Dxz4^∆/∆^*, *Firre*^Xi∆/+^, and *Dxz4^∆/∆^*:*Firre*^Xi∆/+^ (Fig. 8a). We did not observe any “restored” sites on the mutant Xi, when plotting mutant Xi read counts vs. wild-type Xa read counts for peaks that reached at least one half of the wild-type Xa read count (Fig. 8b-d, Fig. S10a). We also compared the Xi read counts for biallelic peaks and observed no changes in *Dxz4^∆/∆^* cells (Fig. 8e), *Firre*^Xi∆/+^ cells, and *Dxz4^∆/∆^*: *Firre*^Xi∆/+^ cells (Fig. 8f, Fig. S10b). Thus, we found no decrease in chromatin accessibility at escapee genes, in contrast to a previous study^22^. Altogether, our results demonstrate that deletion of either *Dxz4*, *Firre*, or both tandem repeats has no impact on chromatin accessibility on the Xi, at either inactive genes or escapees.

**Figure 8:**
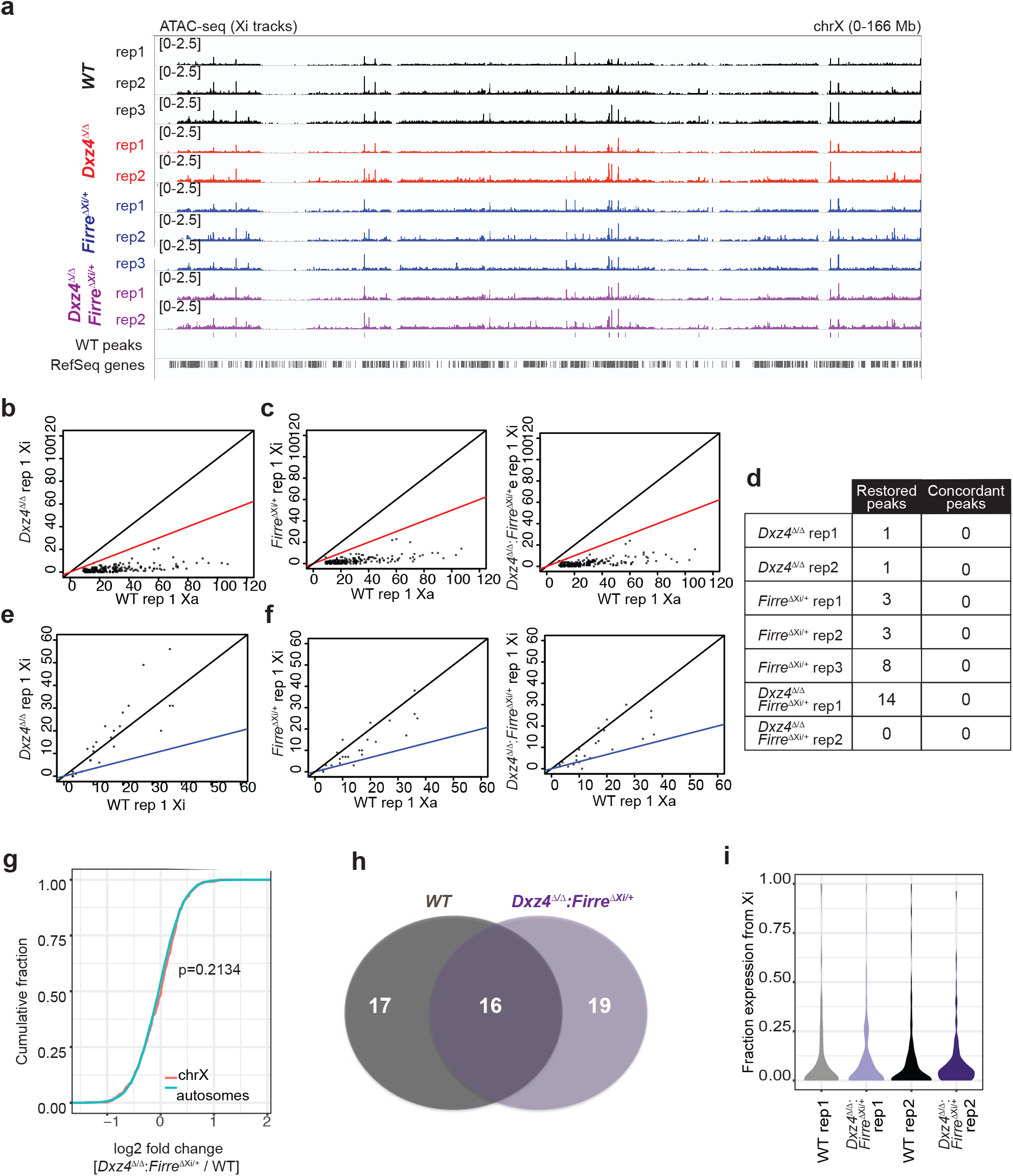
Deletion of *Dxz4*, *Firre*, or both does not affect accessibility or gene silencing on the Xi. (**a**) Comparison of Xi ATAC-seq coverage, WT (black), *Dxz4*^∆/∆^, *Firre*^Xi∆/+^ (blue) and *Dxz4*^∆/∆^:*Firre*^Xi∆/+^ (purple). (**b**) Comparison between WT Xa and *Dxz4*^∆/∆^ Xi ATAC coverage for peaks that are Xa-specific in wild-type. The black line corresponds to a *Dxz4*^∆/∆^ Xi:WT Xa ratio of 1:1, the red line corresponds to a ratio of 1:2. (**c**) Comparison between WT Xa and *Firre*^Xi∆/+^ (left) or *Dxz4*^∆/∆^:*Firre*^Xi∆/+^ (right) Xi ATAC coverage for peaks that are Xa-specific in wild-type. The black line correspond to a *Dxz4*^∆/∆^ Xi:WT Xa ratio of 1:1, the red lines correspond to a ratio of 1:2. (**d**) Number of restored ATAC peaks (Xa-skewed in wild-type but biallelic in deletion) for all deletions. Concordant peaks are peaks reproducibly restored across replicates. (**e**) Comparison between WT Xi and *Dxz4*^∆/∆^ Xi ATAC coverage for peaks that are Xi-specific in wild-type. The black line corresponds to a *Dxz4*^∆/∆^ Xi:WT Xi ratio of 1:1, the blue line corresponds to a ratio of 1:2.(**f**) Comparison between WT Xi and *Firre*^Xi∆/+^ (left) or *Dxz4*^∆/∆^:*Firre*^Xi∆/+^ (right) Xi ATAC coverage for peaks that are Xi-specific in wild-type. The black lines correspond to a *Dxz4*^∆/∆^ Xi:WT Xi ratio of 1:1, the blue lines correspond to a ratio of 1:2. (**g**) CDF of the fold change in gene expression between WT and *Dxz4*^∆/∆^:*FirreXi*^∆/+^ for autosomal genes (teal) and X-linked genes with fpm >1 in all experiments. The Kolmogorov–Smirnov p-value is indicated. (**h**) Overlap between escapees in WT and *Dxz4*^∆/∆^:*Firre*^Xi∆/+^. (**i**) Density plots of the number of genes with a given level of expression from the Xi in two wild-type and two *Dxz4*^∆/∆^:*Firre*^Xi∆/+^ replicate RNA-seq experiments. Grey and black: wild-type, purple: *Dxz4*^∆/∆^:*Firre*^Xi∆/+^.

Finally, we asked whether deleting both *Dxz4* and *Firre* disrupts the pattern of gene silencing or escape on the Xi. We performed allele-specific RNA-seq in wild-type *Tsix^TST^/+* and *Dxz4*^∆/∆^:*Firre*^Xi∆/+^ and looked for changes on either a genome-wide or Xi-scale in the mutant relative to wildtype. Surprisingly, no significant differences were detected on a global or Xi-wide scale (Fig. 8g-i), in contrast to previous deletions of *Dxz4*^22^. Cumulative frequency plots showed balanced X-to-autosomal gene dosages when comparing mutant to wildtype cells (Fig. 8g). The overall number of genes escaping XCI was similar in wild-type and *Dxz4*^∆/∆^:*Firre*^Xi∆/+^, with about half of the escapees being shared between them Fig. 8h). Examination of allelic contributions to overall X-chromosomal expression showed a predominance of Xa expression in both wild-type and *Dxz4*^∆/∆^:*Firre*^Xi∆/+^, with the pattern being highly similar in two biological replicates (Fig. 8i; p > 0.3 for all pairwise comparisons). We conclude that neither *Firre* nor *Dxz4* significantly perturbs Xi silencing and escape from silencing. Thus, the unique superloops of the Xi can be uncoupled from XCI biology.

## DISCUSSION

An outstanding question in genome and nuclear organization is how higher-order chromatin structure regulates gene expression. Here, using the Xi as a model, we have tested the relevance of two higher order structures — superloops and megadomains — for the biology of XCI. While indeed *Dxz4* is required to form megadomains and *Firre* is required for superloops and for full strength of megadomains, we unexpectedly observed that abolishing these structures had no impact whatsoever when assaying a range of XCI phenotypes, including (i) chromosome-wide silencing as determined by RNA-seq, (ii) ability of escapees to avoid inactivation, (iii) subnuclear localization of the Xi, (iv) enrichment of H3K27me3 mark along the Xi, and (v) general chromatin accessibility as measured by ATAC-seq. By analyzing the time course of megadomain and superloop formation, we determined that these structures do not precede Xist spreading or XCI. Instead, they either occur concurrently with Xist spreading and XCI, or they may be a consequence thereof. We also observe that *Dxz4* and *Firre* may work together to strengthen megadomains through superloop formation: Deleting *Firre* weakens intra-megadomain interactions without affecting the strong *Dxz4* border, deleting *Dxz4* abolishes the sharp megadomain border, and deleting both loci has a more severe effect on overall megadomain organization than deleting either singly. The significance of attenuated megadomains is unclear, however, given that there was no perturbation to XCI when either *Dxz4* or *Firre* or both were deleted. Taken together, our data argue that the superstructures are not necessary for XCI biology, at least in the *ex vivo* cellular context.

Our findings therefore beg a number of interesting questions. First, what is the purpose of *Dxz4, Firre,* megadomains, and superloops, and why are they conserved across 80 million years of mammalian radiation? While our observation that *Dxz4* is required for megadomains on the Xi is in agreement with three other studies^22,25,29^, our results are at odds with the previous proposal that *Dxz4* enables genes to escape XCI^22^. Our study also finds no loss of accessibility on ~35 escapee genes when *Dxz4* is deleted on the Xi. The different conclusions may result from either use of different cell types (mESC versus NPCs) or clonal variation. Notably, the previous study observed the effects only in one of four NPC clones and the clone showed an unusually high number of escapees — ~100 escapees, a number that is 2-4 times greater than reported by any other study^50,55,56^. If this NPC clone were an oddity, comparing its expression state to a *Dxz4*-deleted cell line of a different NPC background would lead to the impression that the unusual escapees (which ordinarily do not escape XCI) had become silenced.

Additionally, our data suggest that loss of insulation and formation of the megadomains does not occur before Xist spreading and XCI are completed, and thereby providing further evidence for a decoupling of megadomain structures from epigenetic silencing on the Xi. Indeed, our results are in line with other deletions of *Dxz4*, all of which failed to find an effect on gene silencing after deletion of the megadomains^22,29^. Whatever function *Dxz4* might serve, our study indicates that it is necessary but not sufficient for megadomain formation in post-XCI cells, even when *Xist* is present ectopically together with *Dxz4*. Because we could not derive the transgenic line in a female ES cell background, we do not formally know whether *Dxz4* and *Xist* together might be sufficient during de novo XCI in an ectopic context. However, given that *Dxz4* has no impact on XCI and escape in any measurable way, the sufficiency during the XCI establishment phase seems moot.

*Firre* was also of interest. What is its function and why do superloops form on the Xi via this CTCF-enriched repeat? Although we failed to find an XCI-related function for *Firre* after ablating it on the Xi, we found that *Firre*, *Dxz4*, *Xist*, and *X75* form a conserved network of superloops on the mammalian Xi, in agreement with a previous study^25^. Yet, superloops are dispensable for establishing XCI, since both superloop anchors *Dxz4* and *Firre* can be deleted on the future Xi with no impact on XCI establishment. Perhaps Firre RNA has a role in X-inactivation, as suggested from Firre knockdown experiments implicating Firre RNA in maintaining perinucleolar localization of the of the Xi and enrichment of H3K27me3^55^. We leveraged new Xa- and Xi-specific Firre deletions to test which allele expresses Firre RNA and find distinct isoforms associated with the Xa versus Xi. Quantitative RT-PCR suggests that transcription from the Xa may predominate, though we cannot be certain without knowing all potential isoforms. Deleting *Firre* on either Xi or Xa does not perturb Xi localization, Xist expression or H3K27me3 deposition on the Xi. Thus, neither the *Firre* superloop anchor nor any transcript produced by *Firre* is needed for XCI.

Could megadomains and superloops be default consequences of Xist-mediated attenuation of TADs and compartments? It is important to note that our study follows XCI only in the *ex vivo* cellular context. It is possible that *Dxz4, Firre*, megadomains, and superloops play an important role in long-term maintenance of the Xi and that this role would only be revealed by following mice over their lifespan. It is also possible that these Xi megadomains and superloops are incidental organizational structures with no primary impact on gene regulation. While these structures do not disrupt XCI, other macrostructures do. In particular, the recently identified S1/S2 compartments that are revealed by loss of SMCHD1 function play an essential role during de novo Xi silencing^24^. Why the Xi would be folded in these ways with or without function is unclear. For *Dxz4* and *Firre* superloops, a role in a non-XCI pathway — critical in a whole-organism context and not measurable by our present assays — must also be entertained. Irrespective of function, the megadomains and superloops represent the largest architectural structures identified by Hi-C to date in mammals, and both are clearly unique to the Xi. Their evolutionary conservation across 80 million years suggests that the superstructures likely persist for reasons that will only become clear with further study.

## CONTRIBUTIONS

J.E.F. and J.T.L. designed research, analyzed data, and wrote the manuscript. J.E.F. deleted *Dxz4* and *Firre*, performed all Hi-C, RNA-seq microscopy experiments and all bioinformatics except for 4C analysis. S.F.P. designed, performed, and analyzed 4C experiments. A.J.K. performed the Hi-C^2^ experiments. T.J. performed ATAC-seq experiments.

**Data availability:** All sequencing data that support the findings of this study have been deposited in the National Center for Biotechnology Information GEO repository under accession GSE116649.

## ACKNOWLEDGEMENTS

We thank Chen-Yu Wang and Eric Aeby for a careful reading of the manuscript. This work was supported by NSF GRFP grants (J.E.F., A.J.K.), a Herchel Smith Fellowship (A.J.K.), and an NIH grant R01-GM090278 (J.T.L.). T.J. is an EMBO postdoctoral fellow (EMBO ALTF 1313-2015). J.T.L. is an investigator of the Howard Hughes Medical Institute.

## SUPPLEMENTAL FIGURES

**Figure S1:**
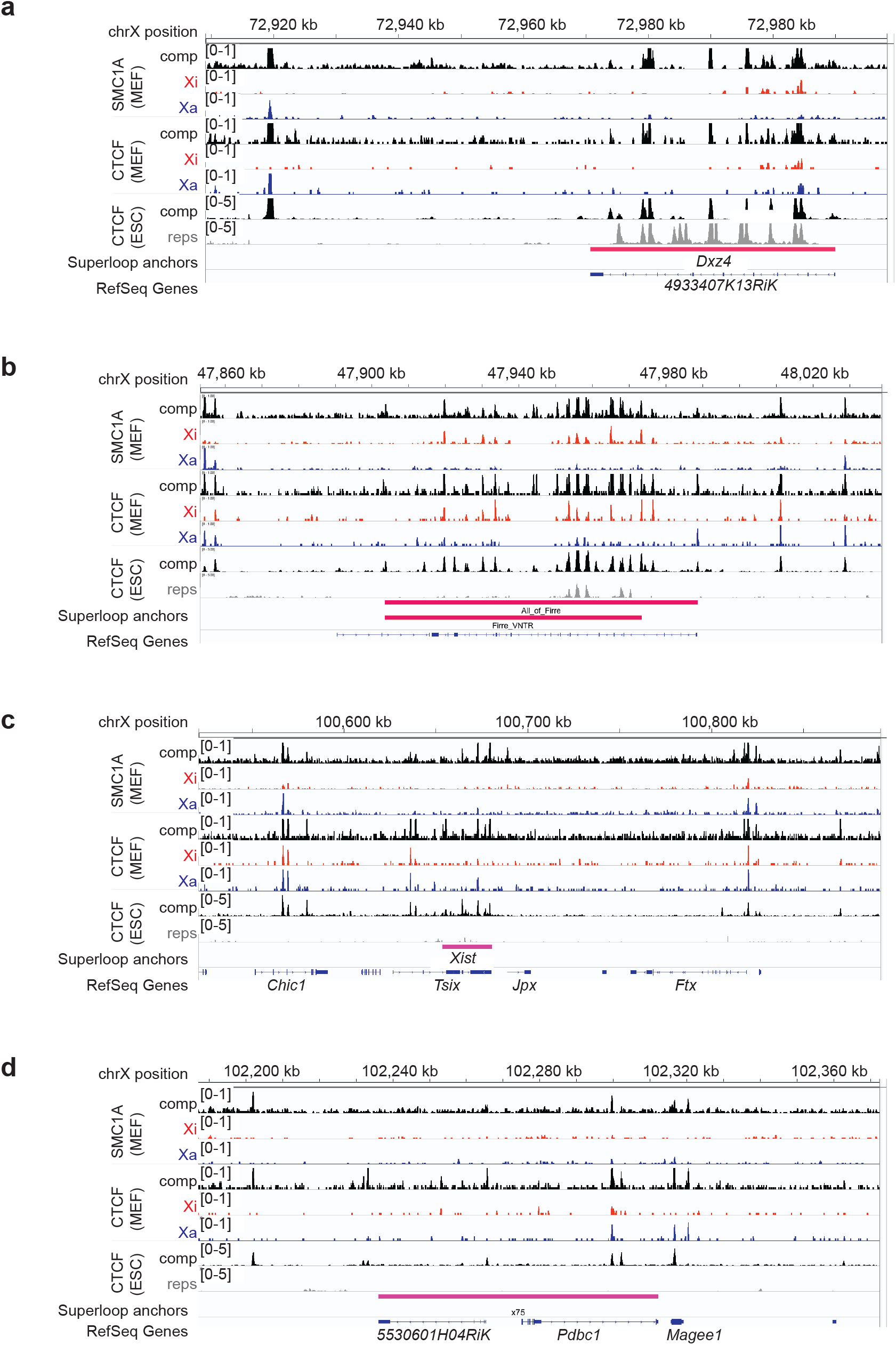
Superloop anchors are tandem repeats that bind CTCF and SMC1A on the mouse Xi. SMC1A and CTCF ChIP-seq coverage across the superloop anchors in mouse (**a**) Dxz4, (**b**) Firre, (**c**) The X-Inactivation Center, and (**d**) x75. From top to bottom within a panel: SMC1A ChIP-seq in MEF, black=comp (all unique reads including neutral, cas, and mus), blue=Xa, red=Xi, CTCF ChIP-seq in MEF (same color scheme), CTCF ChIP-seq in mESCs, black=comp, grey=repetitive alignments.

**Figure S2:**
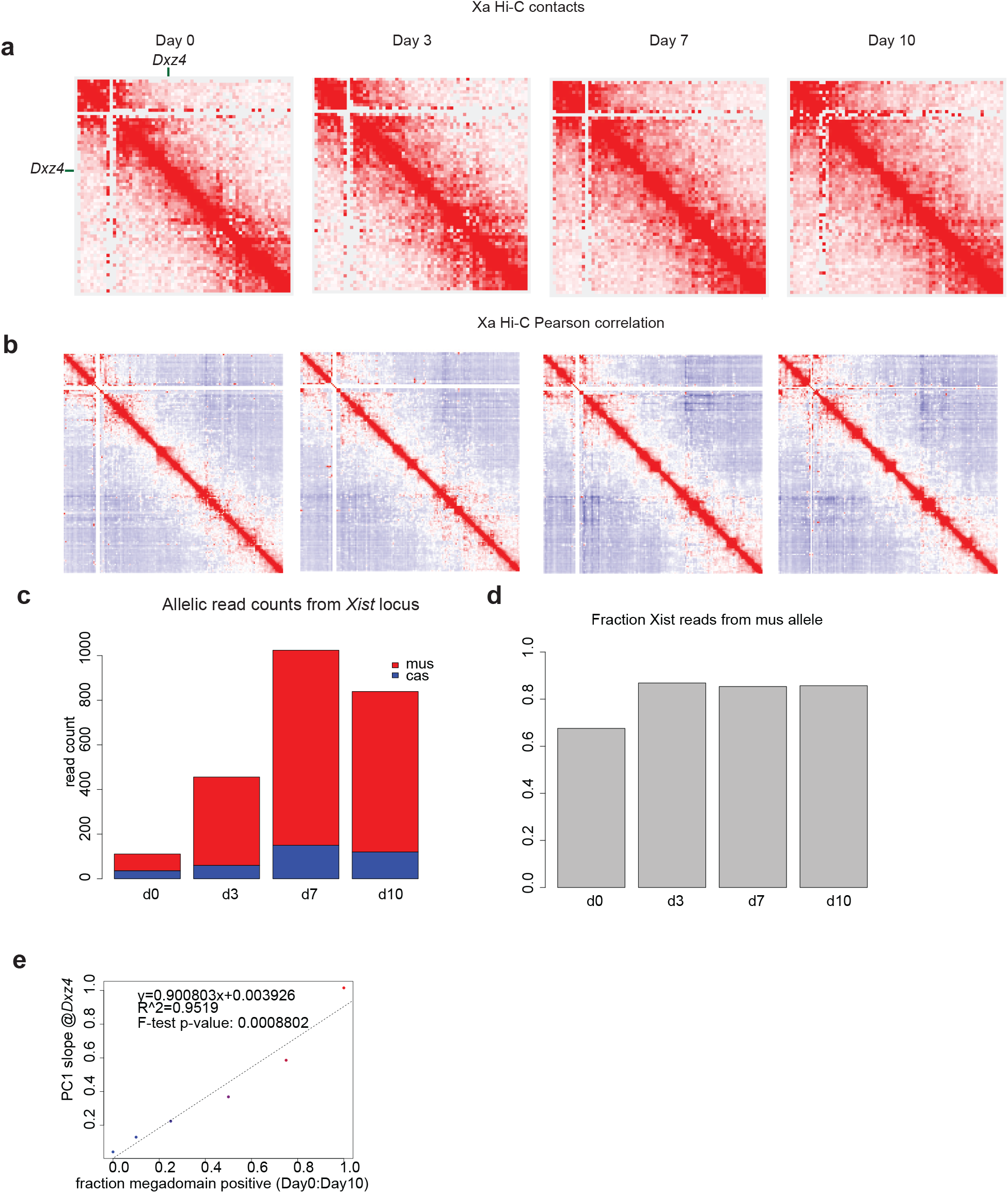
Megadomains are not present on the Xa and allelic RNA-seq shows Xist upregulation is highly skewed to Xi. (**a**) KR-normalized Hi-C matrices on future Xa (cas) on days 0, 3, 7,10 of differentiation (2.5 Mb resolution). (**b**) Pearson correlation of Hi-C matrix on future Xa (cas) on days 0, 3, 7, 10 of differentiation (1Mb resolution). **(c)** Numbers of Xist reads expressed from the mus (red) or cas (blue) during differentiation. **(d)** Fraction of allelic Xist reads expressed from X^mus^ (carrying the *Tsix^TST^* allele) during differentiation, in two biological replicates. (**e**) Slope of the PC1 score curve at *Dxz4* for Hi-Cs generated with varying ratios of d10:d0 reads. Dashed line represents linear best fit; linear model parameters are included.

**Figure S3:**
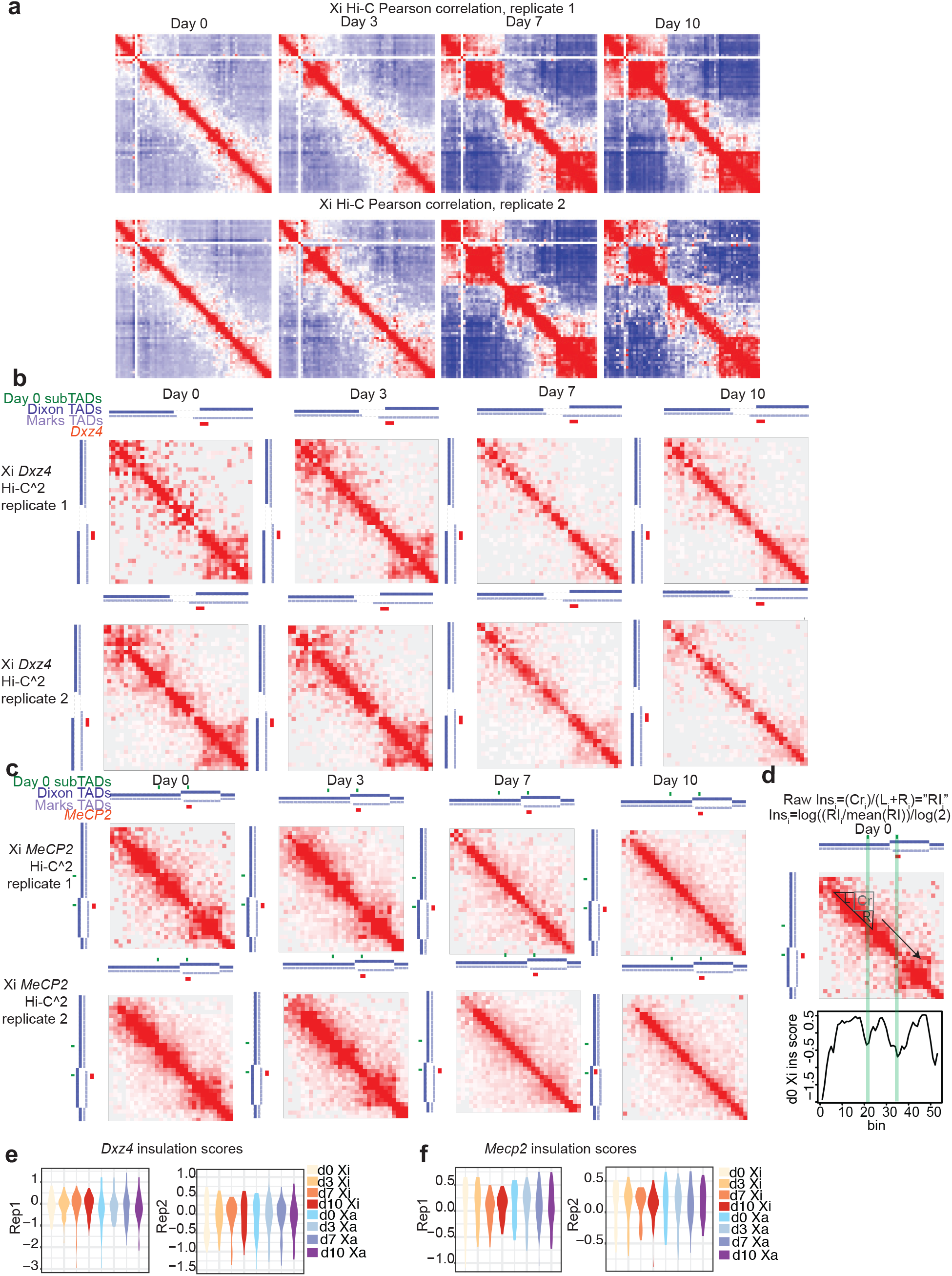
Biological replicates of the time course Hi-C and Hi-C^2^ experiments. **(a)** Pearson correlation matricies at 2.5 Mb resolution for the Xi (right) days 0, 3, 7, 10 of differentiation in two biological replicate Hi-C experiments. (**b**) Hi-C^2^ contact maps around *Dxz4* (mm9 coordinates chrX:71,832,976-73,511,687) and on the Xi on days 0, 3, 7, 10 of differentiation (50 kb resolution) in two biological replicates. (**c**) Hi-C^2^ contact maps around *Mecp2* (mm9 coordinates chrX:70,370,161-71,832,975) and on the future Xi on days 0, 3, 7, 10 of differentiation (50 kb resolution) in two biological replicates. In **(b)** and **(c)**, Green bars indicate positions of domain borders determined from 25 kb d0 comp Hi-C^2^ matrices; dark blue track shows Dixon et al. TAD calls in mESCs, light blue track shows Marks et al. TAD calls in mESCs, red bars indicate positions of either *Dxz4* or *Mecp2.* **(d)** Conceptual diagram of insulation score. Top: formula for calculating insulation score at region i. R_i_ refers to the sum of interactions within the region to the right of i, L_i_ refers to the sum of interactions within the region to the left of i, and Cr_i_ represents the sum of interactions that “cross over” i. Middle: diagram of the window used to calculate insulation score for an example (non-border) i in the day 0 Xi *Mecp2* contact map. Bottom: plot of insulation score across the day 0 Xi *Mecp2* region. The shaded green bars indicate the positions of the borders of the Mecp2 sub-TADs across all diagrams. **(e, f)** Violin plots showing the distributions of insulation scores across the *Dxz4* region (**e**) and *Mecp2* region **(f)** in both biological replicates. Note: to generate violin plots and evaluate the significance of differences in variance between timepoints we excluded the 6 bins on each edge of the Hi-C^2^ region because the regions needed to calculate insulation score fall partly outside the Hi-C^2^ region and have far lower read counts than sequences targeted by the capture probes.

**Figure S4:**
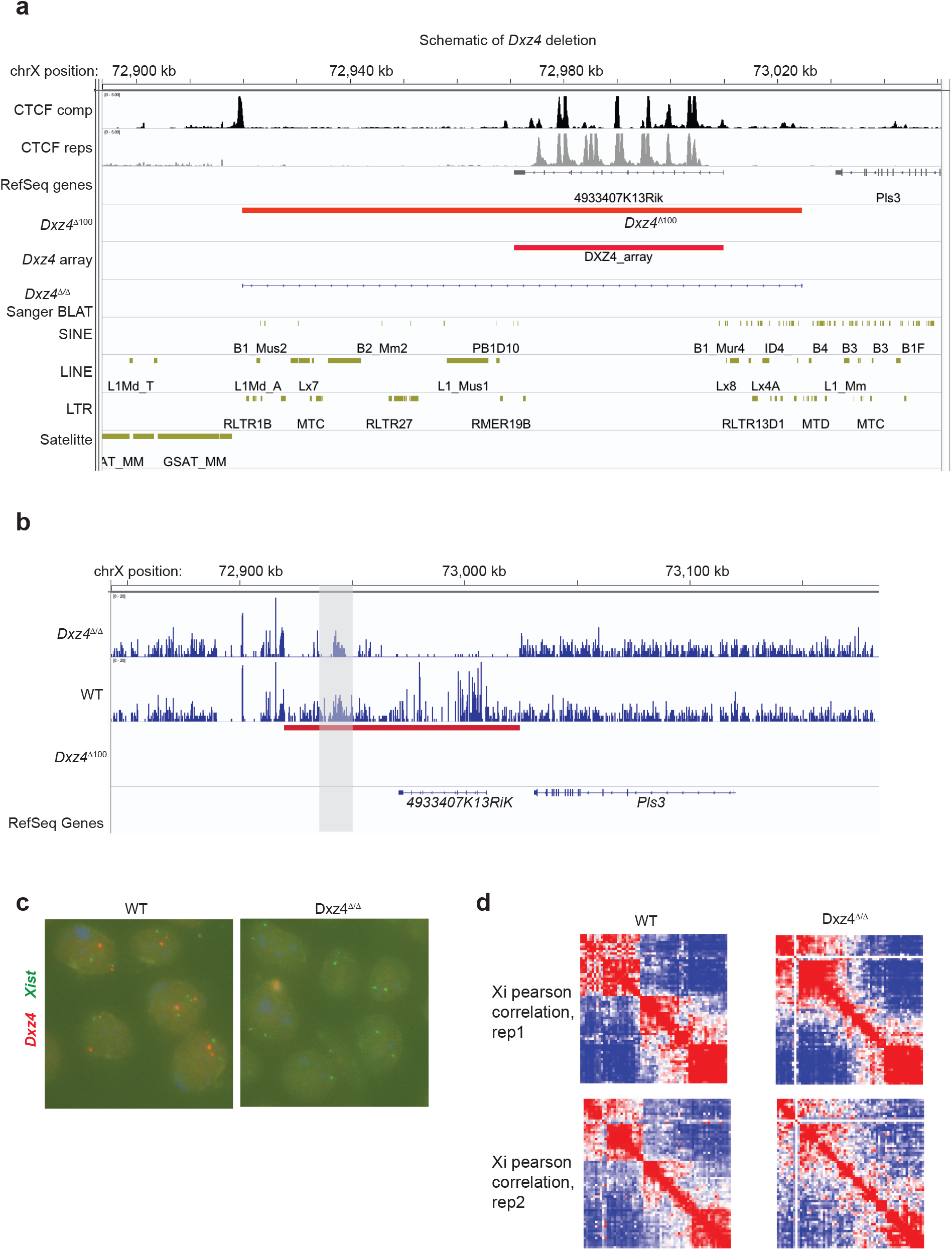
Validation of *Dxz4* deletion and replicate Hi-C. (a) Scheme of the *Dxz4* deletion. The large red bar shows the region deleted in this study; the smaller red bar shows the coordinates of the *Dxz4* tandem repeat array. Interspersed repeats are shown in gold, and CTCF ChIP-seq in mESCs is shown in black for all unique reads and grey for all repetitive reads. The “∆Dxz4 Sanger BLAT” track is the BLAT result for Sanger sequencing result from the PCR product generated with primers flanking the deleted region. (**b**) Hi-C coverage over the *Dxz4* deletion region for *Dxz4*^∆/∆^ clone E5 (top) and WT (bottom). The near-absence of reads in *Dxz4*^∆/∆^ over the deletion implies a biallelic deletion. Note: we observed reads in *Dxz4*^∆/∆^ within the left end of the deletion (grey box), roughly chrX:72,940,913-72,948,483. BLAT analysis of this region indicates that it is repeated several times elsewhere in the genome (data not shown), thus reads within this region are to be expected even in *Dxz4*^∆/∆^. (**c**) DNA FISH in WT (left) and *Dxz4*^∆/∆^ (right) using either *Dxz4* (red) or *Xist* (green) probes. (**d**) Pearson correlation matricies at 2.5 Mb resolution for the wild-type (left) and *Dxz4*^∆/∆^ Xi (right) in two biological replicate Hi-C experiments. Note: the WT control Hi-C in replicate 2 in this figure also serves as the replicate 2 day 10 timepoint in FigS3a, thus this Pearson correlation matrix is present in both FigS3a and this figure.

**Figure S5:**
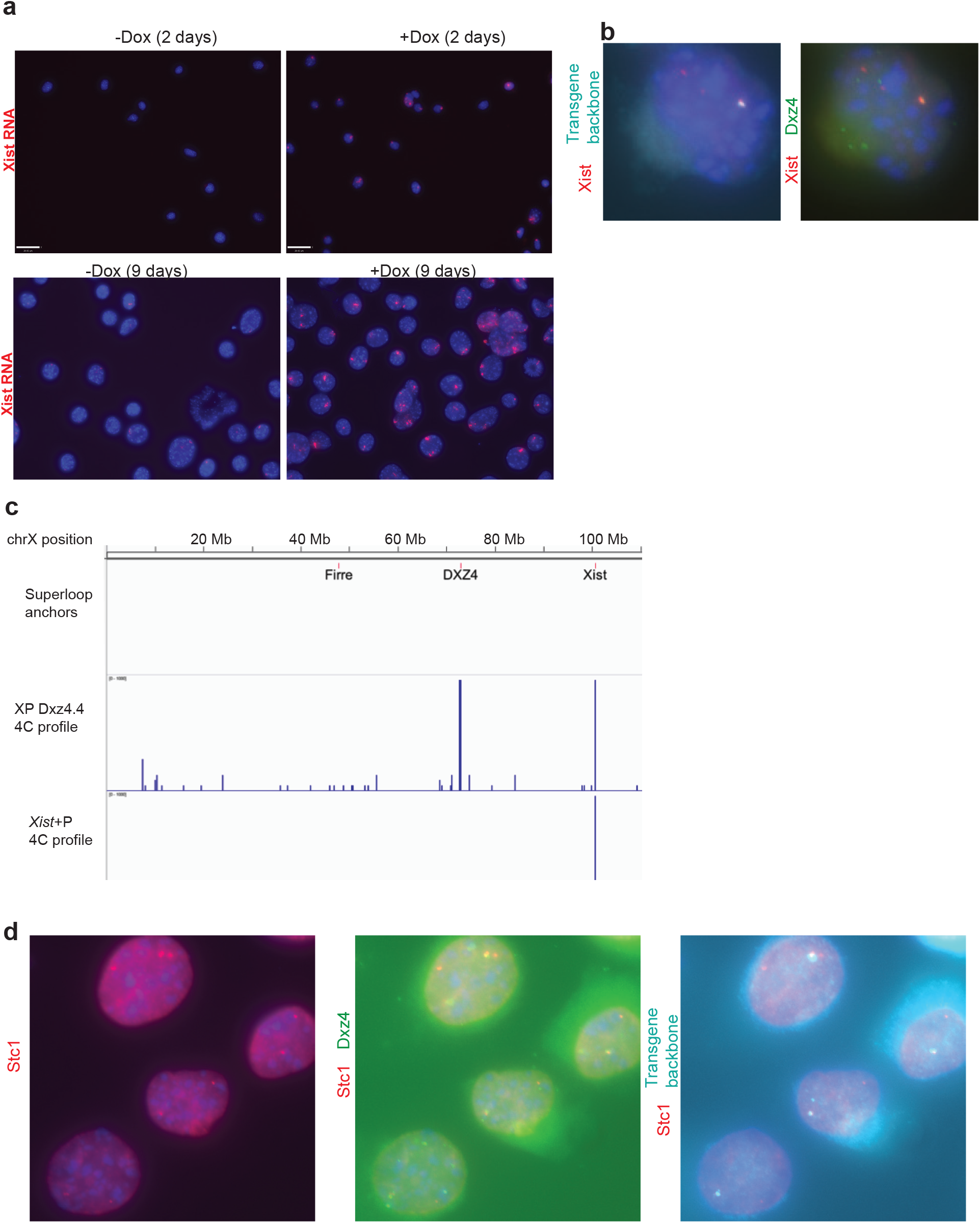
Generation of a *Xist+Dxz4* transgene. (**a**) Xist RNA FISH in *Xist+Dxz4* Tg cells (clone XPDxz4.4) +/-dox after either 2 days of induction (top) or 9 days of induction (bottom panels). (**b**) Co-localization between transgenic *Xist* and *Dxz4*. Left, DNA FISH for the transgene’s P1 backbone (cyan)+*Xist* (red). Right: DNA FISH for *Xist* (red) and Dxz4 (green). (**c**) 4C contact profiles in XPDxz4.4 or a separate *Xist*-only transgenic cell line (XY X+P) on chrX from a viewpoint within the backbone of the *Xist* construction. The positions of *Firre, Dxz4* and *Xist* are indicated on the X-chromosome. (**d**) Co-localization between *Xist* and *Dxz4* and the candidate insertion region (*Stc1*, chr14) obtained from 4C. Left: DNA FISH for *Stc1* (red). Middle: overlap between DNA FISH for *Stc1* (red) and *Dxz4* (green). Right: overlap between DNA FISH for *Stc1* (red) and the Xist transgene’s P1 backbone (cyan).

**Figure S6:**
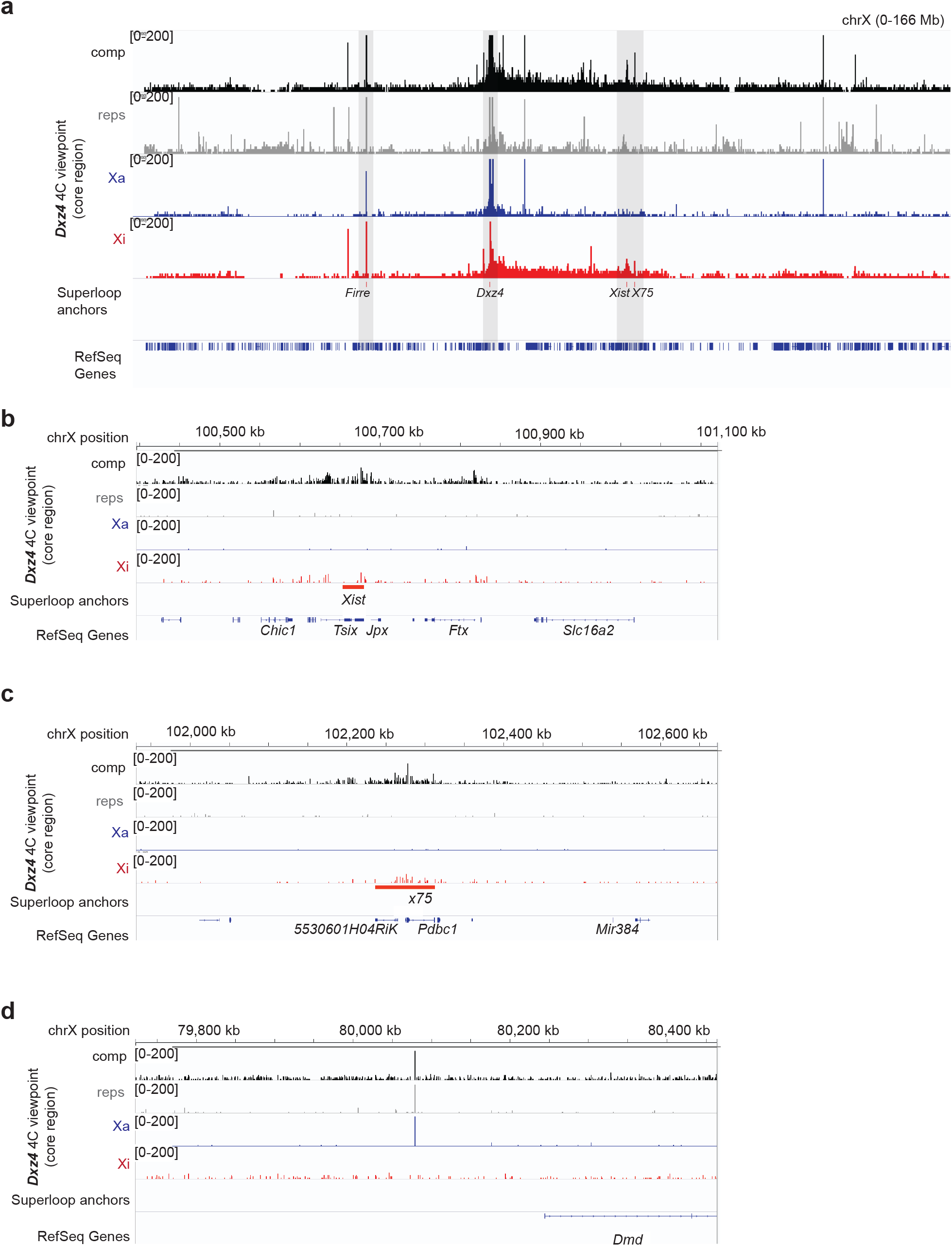
A conserved network of superloop interactions involving *Dxz4* on the mouse Xi. (**a**) 4C coverage from the *Dxz4* viewpoint over the whole X in mus Xi MEFs. Black=comp, grey=reps, blue=Xa, red=Xi. The positions of putative superloop anchors *Firre, Dxz4, Xist* and *x75* are shown. (**b**) 4C coverage from the *Dxz4* viewpoint over the *Xist* region. (**c**) 4C coverage from the *Dxz4* viewpoint over the region homologous to human x75 in mouse. (**d**) 4C coverage from the *Dxz4* viepoint over a putative singleton artifact near *Dmd*. Many of the other large peaks on the X or elsewhere are “singletons”, amplified sequences likely due to mispriming, this is an example of a singleton.

**Figure S7:**
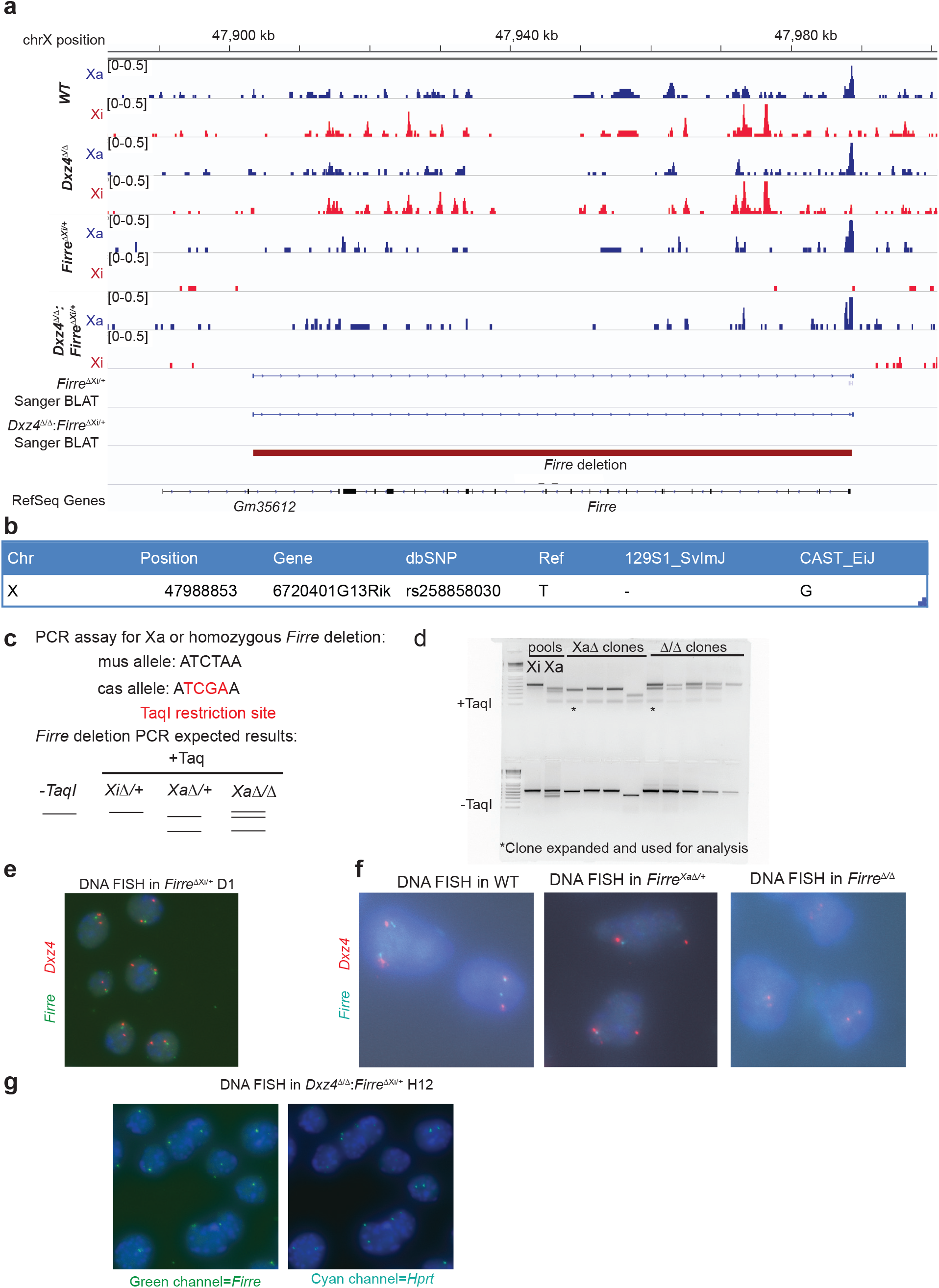
Validation of *Firre* deletions. (**a**) Allele-specific ATAC-seq coverage over *Firre*. The coordinates of the *Firre* deletion are shown by the dark red bar, and the two “Sanger BLAT” tracks represent the coordinates of the BLAT alignment of the PCR products flanking Firre generated from *Firre*^Xi∆/+^ clone D1 and *Dxz4^∆/∆^:Firre^Xi∆/+^* clone H12. **(b)** SNP information from the Mouse Genomes Project (cite Sanger) for the SNP that falls within the *Firre* deletion amplicon. **(c)** Scheme of restriction assay to determine which allele of *Firre* is deleted. **(d)** Results of restriction assay in Xa- and Xi- Firre targeted pools and *Firre*^Xa∆/+^ and *Firre*^∆/∆^ candidate clones (**e**) DNA FISH in *Firre*^Xi∆/+^ clone D1 for *Dxz4* (red), and *Firre* (green). **(f)** DNA FISH for *Dxz4* (red) and *Firre* (cyan) in WT, *Firre*^Xa∆/+^ and *Firre*^∆/∆^ clones. In *Firre*^Xi∆/+^ and *Firre*^Xa∆/+^ most cells exhibit one *Firre* spot but two *Dxz4* spots, consistent with a heterozygous *Firre* deletion, whereas *Firre*^∆/∆^ does not exhibit Firre spots, consistent with a homozygous *Firre* deletion. (**g**) DNA FISH in *Dxz4*^∆/∆^:*Firre*^Xi∆/+^. Left, *Firre* (green). Right, X-linked gene *Hprt* (Cyan). Most cells exhibit one *Firre* spot but two *Hprt* spots, consistent with a heterozygous *Firre* deletion.

**Figure S8:**
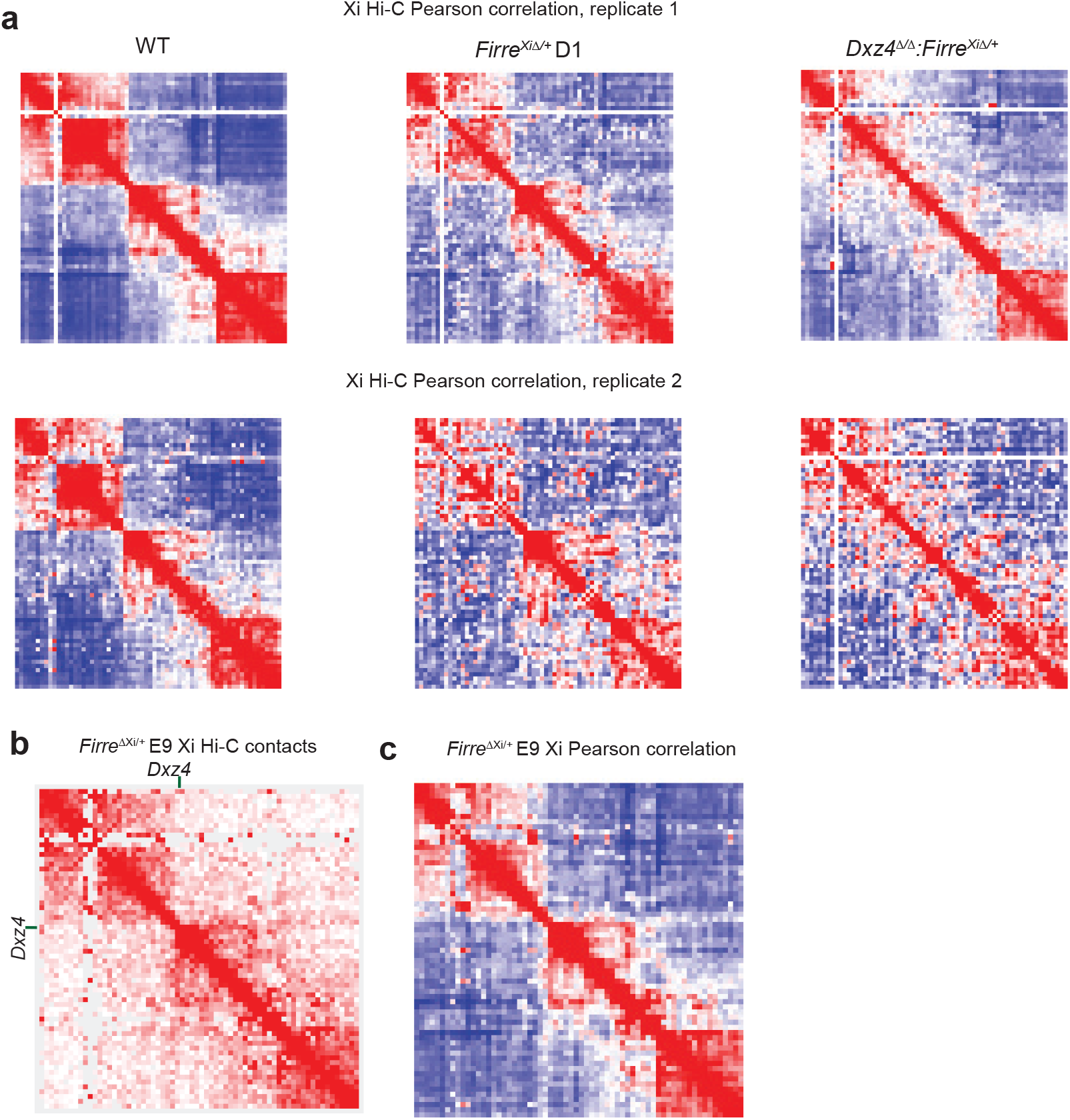
Replicate Hi-Cs in *Firre* deletions. **(a)** Pearson correlation matrices at 2.5 Mb resolution for the wild-type (left) and *Firre*^*Xi*∆/+^ (middle, clone D1) and *Dxz4*^∆/∆^:*Firre*^Xi∆/+^ Xi (right) in two biological replicate Hi-C experiments. Note: the WT control Hi-C in replicates 1 and 2 in this figure also serve as the replicates 1 and 2 day 10 timepoints in FigS3a, thus these Pearson correlation matrices are present in both FigS2a and the current figure. **(b)** KR-normalized Hi-C matrix for the independently derived *Firre*^Xi∆/+^ clone (E9) at 2.5 Mb resolution. **(c)** Pearson correlation matrix for the independently derived *Firre*^Xi∆/+^ clone (E9) at 2.5 Mb resolution.

**Figure S9:**
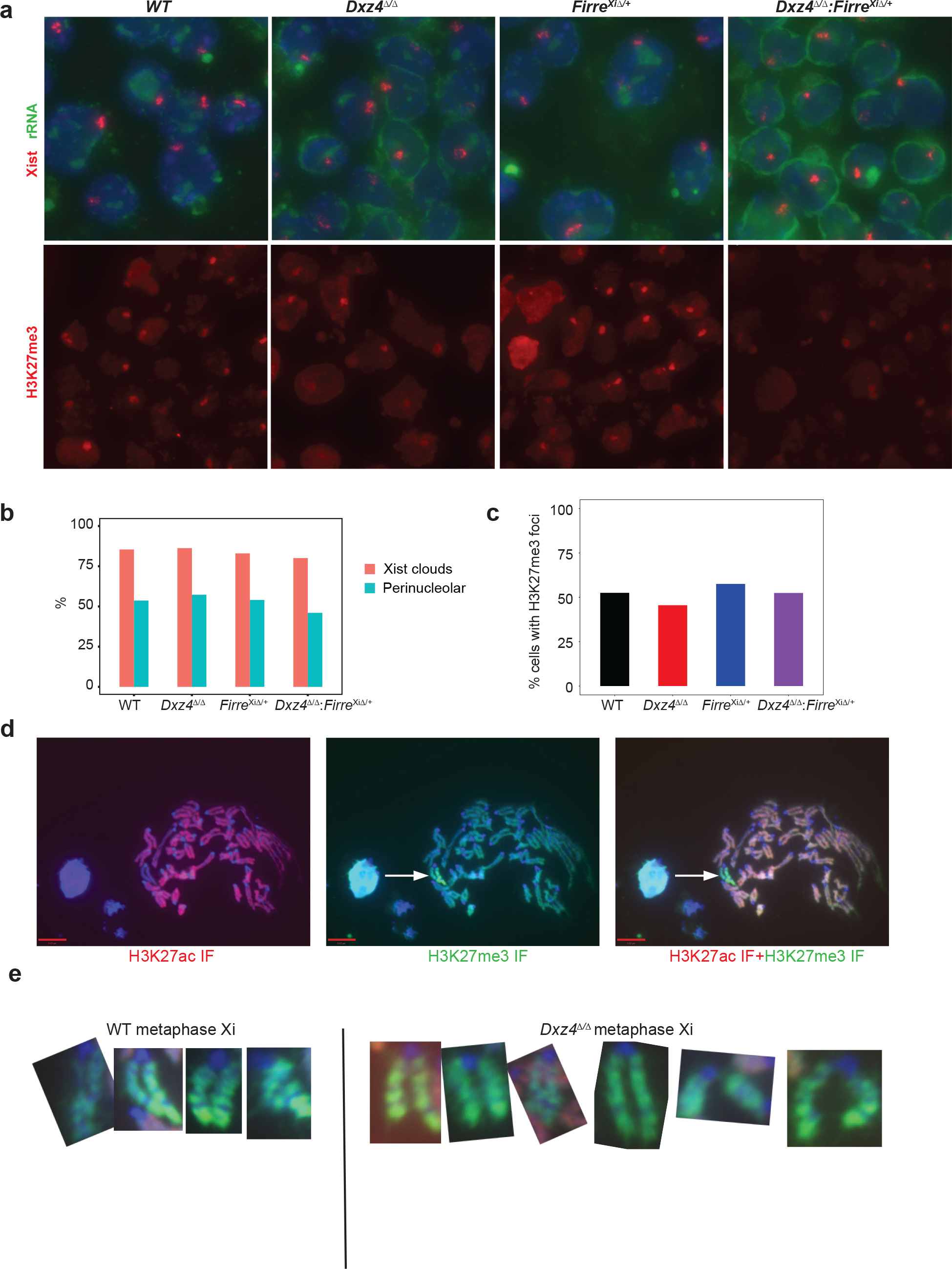
Further analysis of Xist localization and H3K27me3 foci in *Dxz4* and *Firre* deletions; metaphase distribution of H3K27me3 in *Dxz4*^∆/∆^. (**a**) Xist RNA FISH (red) combined with rRNA FISH (green) (top), H3K27me3 IF (red, bottom) in WT and *Dxz4*^∆/∆^ *Firre*^Xi∆/+^ and *Dxz4*^∆/∆^:*Firre*^Xi∆/+^ cells on day 10. (**b**) Fraction of cells with Xist clouds (red) and fraction of Xist clouds in the perinucleolar space (cyan) on day 10. (**c**) Fraction of cells with an H3K27me3 focus. **(d)** Immunoflourescence for H3K27ac (red, left) or H3K27me3 (green, middle) on metaphase spreads from WT cells after 10 days of differentiation. The Xi is readily detectable as the one chromosome in the spread depleted of H3K27ac and enriched in H3K27me3 (merge, right). **(e)** Coating of H3K27me3 on several metaphase inactive Xs in WT (left) or *Dxz4*^∆/∆^ (right) on day 10.

**Figure S10:**
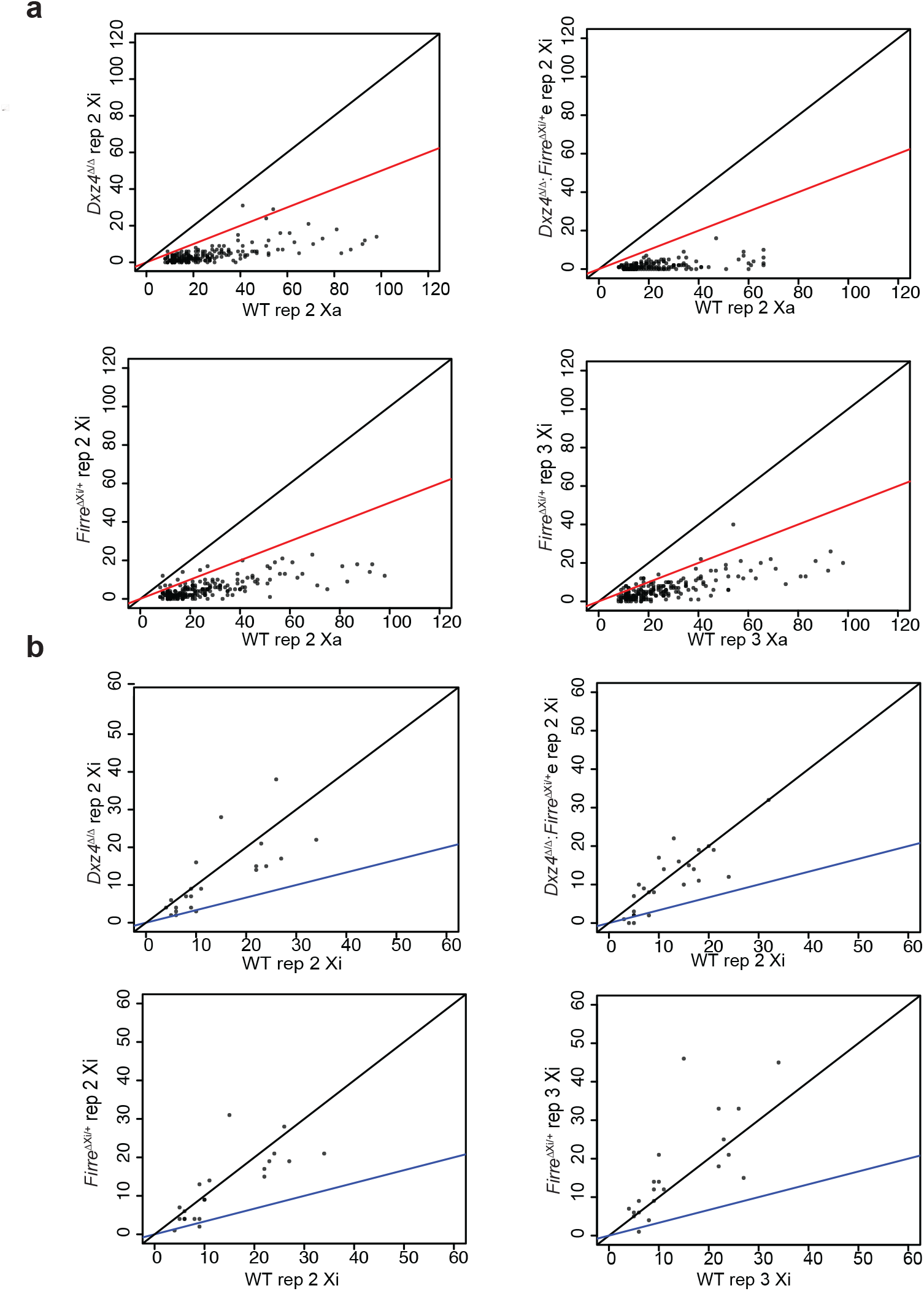
Replicate ATAC-seq analysis in *Firre*Xi^∆/+^ and *Dxz4*^∆/∆^:*Firre*^Xi∆/+^. (**a**) Comparison between WT Xa and mutant Xi for ATAC coverage for peaks that are Xa-specific in wild-type for replicate ATAC-seqs. Top left: *Dxz4*^∆/∆^ Xi vs WT Xa, top right: *Dxz4*^∆/∆^:*Firre*^Xi∆/+^ Xi vs WT Xa, Bottom left: *Firre*^Xi∆/+^ Xi vs WT Xa, replicate 2, Bottom left: Firre^Xi∆/+^ Xi vs WT Xa, replicate 3. Black lines corresponds to a *Dxz4*^∆/∆^ Xi:WT Xa ratio of 1:1, red lines corresponds to a ratio of 1:2. (**b**) Comparison between WT Xa and mutant Xi ATAC coverage for peaks that are Xi-specific in wild-type for replicate ATAC-seqs. Top left: *Dxz4*^∆/∆^ Xi vs WT Xi, top right: *Dxz4*^∆/∆^:*FirreXi*^∆/+^ Xi vs WT Xi, Bottom left: *Firre*^Xi∆/+^ Xi vs WT Xi, replicate 2, Bottom left: *Firre*^Xi∆/+^ Xi vs WT Xi, replicate 3. Black lines correspond to a *Dxz4*^∆/∆^ Xi:WT Xi ratio of 1:1, blue lines correspond to a ratio of 1:2.

## SUPPLEMENTAL INFORMATION

### MATERIALS & METHODS

#### Cell lines and growth conditions

ES cells were grown in regular ES+LIF medium (500 ml DMEM with the addition of 1 ml of β-mercaptoethanol, 6 ml of MEM NEAA, 25 ml of 7.5% NaHCO_3_, 6 ml of GlutaMAX-1, 15 ml of 1M HEPES, 90 ml of FBS, 300 μl of LIF, 6 ml of PEN/STREP) on irradiated feeders. To differentiate mESCs and allow them to undergo X-inactivation, mESCs were harvested by trypsinization and quenched in ES medium without LIF. Feeders were removed by adding the cell suspension to tissue culture plates for 45 minutes at 37°C. Differentiating embryoid bodies were cultured for 4 days on low-adherence plates in ES medium without LIF. On the 4^th^ day, the embryoid bodies were plated onto gelatinized tissue culture plates and allowed to attach. ES cells for experiments were harvested by extensive trypsinization to detach them from the plates. Unless otherwise noted, all experiments were performed after 10 days of differentiation.

Fibroblasts were grown on un-gelatinized tissue culture plates in MEF media (500 ml DMEM with the addition of 1 ml of β-mercaptoethanol, 6 ml of MEM NEAA, 25 ml of 7.5% NaHCO_3_, 6 ml of GlutaMAX-1, 15 ml of 1M HEPES, 60 ml of FBS, 300 μl of LIF, 6 ml of PEN/STREP).

#### Generation of *Dxz4* and *Firre* deletion cell lines

Oligos encoding gRNAs flanking either *Dxz4* or *Firre* were cloned into wild-type Cas9+GFP plasmid PX458^1^ by first linearizing the plasmid with BbsI, purifying the plasmid using the Qiagen PCR Purification Kit, then ligating annealed and phosphorylated oligos into the plasmid using T4 DNA ligase for 1 hour at room temperature, then transforming into OneShot Top10 chemically competent *E. coli*. gRNA plasmid DNA was prepared using the Qiagen Miniprep Kit and gRNA sequences were verified by Sanger Sequencing. To delete *Dxz4* or *Firre*, pairs of gRNAs flanking either *Dxz4* or *Firre* were transfected into mESCs using Lipofectamine LTX. Briefly, for each transfection, 50 uL of OptiMEM media was added to 2.5 uL LTX reagent and 50 uL OptiMEM was added to 0.5 uL PLUS reagent. The OptiMEM+PLUS mix was added to a mixture of 250 ng of each gRNA, then the OptiMEM+LTX mix was added and incubated at room temperature for 5 minutes to generate the transfection mixture. Meanwhile, 2×10^5^ mESCs were harvested by trypsinization and brought to a volume of 900 uL ES+LIF media. Once the transfection mixture was ready, the mESCs were layered dropwise on top of it and allowed to incubate for 20 minutes at room temperature. Following incubation, the entire transfection mixture was added to one well of a 12-well dish containing feeders and 1 mL ES+LIF media. The transfected cells were allowed to grow for 16-48 hours.

To screen for Cas9-transfected cells, the transfected cells were harvested with trypsin, washed 2X in PBS and resuspended in 300 uL FACS media (1X Leibowitz’s+5% FBS) and passed through a cell strainer. The GFP-positive cells were isolated by FACS selection and plated on 10 cm feeder plates (~2000-10,000 GFP+ cells/plate). The FACS-sorted GFP positive cells were allowed to grow into large colonies, typically after about 6-8 days of growth. 192 colonies were manually picked and transferred to 96 well plates covered in feeders. Once the 96 well plates were nearly confluent, they were passaged onto 3 new gelatinized plates (no feeders). Freezing media (MEF media+ final concentration 10% DMSO) was added to two plates and they were left at −80°C for storage. The third plate was grown until most wells were fully confluent.

We used a PCR screen to identify *Dxz4* or *Firre* deletion clones. Genomic DNA was prepared from the colonies by incubating them overnight in Laird buffer+proteinase K (50 uL buffer per well) at 55°C. The genomic DNA was transferred to a new 96 well plate and diluted it 1:10 in H2O, then heated at 95°C for 10 minutes to denature it and inactivate the proteinase K. Next, PCR reactions using primers flanking *Dxz4* or *Firre* were prepared in 96 well plates using 20 uL PCR mix+2 uL denature genomic DNA. 40 cycles of amplication were used, and the PCR reactions were run on 2% agarose gels and visualized by ethidium bromide staining. Deletion clones were identified by PCR reactions that produced a band at the expected size. Deletion clones were thawed onto 12-well plates with feeders, and deletions were verified by Sanger sequencing the PCR product, performing DNA FISH using a fosmid probe entirely within the deleted region, and examining reads over the deleted regions from our genomics experiments.

To generate Xa-specific and homozygous *Firre* deletions, we employed a restriction assay to determine whether clones carried a deletion on the Xa or Xi (or both). We took advantage of a cas- (Xa-) specific polymorphism that creates a new TaqI restriction site within the Firre deletion PCR product to screen clones for deletions on particular alleles. We performed PCR amplification as before, but then added 30 uL of 1X Cutsmart buffer+10U TaqI (NEB) to each PCR reaction, then incubated the reactions at 65°C for 45 minutes before running the reactions on a 2% agarose gel.

#### Preparation of high molecular weight DNA

Briefly, 500 mL cultures of *E. coli* containing either Xist+P or RP23-161K4 were grown and spun at 4000 rpm for 15 minutes. Alkaline lysis was performed by re-supsending in 20 mL Buffer 1, aliquoting the cell suspension into two Oak Ridge polypropylene centrifuge tubes, adding 10 mL of Buffer 2 to each tube and inverting 20 times to mix, then adding 12 mL buffer 3 and inverting 20 times and incubating on ice for 5-10 minutes. Protein and genomic contaminants were removed by centrifugation at 10,000 rpm in a JA-20 rotor at 4°C. DNA was precipitated by adding 35 mL isopropanol to 15 mL centrifuged lysate in a 50 mL Falcon tube, incubating 20 minutes at room temperature, then spinning at 3500 rcf for 20 minutes at 4°C. Pellets were resuspended in 500 uL TE+1%SDS and 15 uL 20 mg/mL Proteinase K was added and the DNA mixture was incubated at 55°C for 1.5 hours to remove protein contaminants. The DNA was phenol:choloroform extracted by adding phenol:cholorform:isoamyl alchohol and shaking by hand for 20 seconds, then the DNA was precipitated with 40 uL 3M NaOAc and 1 mL isopropanol per 500 uL DNA mixture for 10 minutes at −20°C. At this time, the precipitated DNA formed a stringy white mass, the excess liquid was removed from this mass and 1 mL 70% ethanol was added to the DNA. The DNA was centrifuged for 5 minutes at 16300 g, the supernatant removed and 1 mL 70% ethanol was added to the pellet and the pellet was spun again for 5 minutes at 16300 g. Supernatant was removed and excess ethanol was allowed to evaporate for 5 minutes, then the DNA pellet was resuspended in 200 uL 10 mM Tris by gentle pipetting with a cut tip.

#### Generation of a *Xist*+*Dxz4* transgene

To generate an autosomal *Xist*+Dxz4 transgene, we co-transfected a doxycycline-inducible *Xist* construct and a BAC containing mouse *Dxz4* into male fibroblasts containing rtTA^2^. First, we prepared DNA from our “Xist+P” construct and the *Dxz4-*containing BAC RP23-161K4 using a custom high-molecular weight purification protocol. DNA for transfection was only used if it gave the expected digest pattern with either XhoI+BamHI for *Xist*+P of XhoI for RP23-161K4 and was not excessively smeared. To co-transfect *Xist*+P and RP23-161K4 into fibroblasts, 2×10^6^ fibroblasts were harvested, washed twice in 1X PBS and resuspended in 700 uL 1X PBS. We then added 5 ug *Xist*+P and 20 ug RP23-161K4 to the cell suspension and electroporated in a 1 mm cuvette at 200V, 1050 uF using a Bio-RAD Xcell GenePulser electroporation system. Electroporated cells were plated onto 3 10 cm dishes in MEF media made with tet-free FBS and grown for one day. The *Xist*+P construct contains a hygromycin selectable marker, and to select for *Xist* transgenes, starting one day after transfection, we added 200 ug/mL hygromycin to the media and changed the media every day for 10 days. Once colonies were grown, we manually picked them and transferred them to 96-well plate. We only obtained about 10 hygromycin resistant colonies. Once confluent, we split colonies onto 3 wells of a 24-well plate. One well was kept for maintenance, the other two were used for screening.

To screen for transgenic lines with both inducible *Xist* and *Dxz4* inserted at the same ectopic site, we used the following strategy. We induced each clone with 1000 ug/mL doxycycline overnight and performed Xist RNA FISH to test if *Xist* could be induced. We also performed Xist RNA FISH in the same clones without dox induction to ensure *Xist* expression is insducible. We kept clones that could induce robust Xist RNA FISH clouds. We then used DNA FISH to check whether the *Xist*+P construct inserted at the same site as the *Dxz4*-containing BAC. We simultaneously performed DNA FISH using an *Xist* probe, a probe within the *Xist* construct backbone, and a fosmid probe against *Dxz4*. We obtained one clone where all 3 probes co-localize at one spot, indicating co-insertion of *Xist* and *Dxz4* into an autosome. We then used 4C to localize the candidate insertion site into *Stc1* on chr14. We then performed DNA FISH using a fosmid probe overlapping *Stc1* combined with a *Dxz4* fosmid and a probe overlapping the backbone of the *Xist* transgenic construct to confirm co-localization of *Xist*, *Dxz4* and *Stc1* at one spot.

#### DNA FISH

BAC or fosmid DNA was prepared using the high molecular weight DNA preparation procedure. Probes were labeled using the Roche Nick Translation kit. 75,000-150,000 cells were cytospun onto slides for 5 minutes at 1000 rpm. Cells were pre-extracted and fixed by passing the slides through CSK-T for 3 minutes at 4°C, CSK for 3 minutes at 4°C, 1X PBS+4% formaldehyde for 10 minutes at room temperature. RNA was removed by digestion with 0.1 mg/mL RnaseA in 1X PBS for 1 hr at 37 degrees. Slides were dehydrated by passage through 70%, 90%, 100% ethanol for two minutes at each concentration, then allowed to dry. Probe was added to hybridization mix (50% formamide, 2X SSC, 10% dextran sulfate, 0.1 mg/mL mouse Cot-1 DNA) and added directly to the slides. Slides were denatured at 92°C for 10 minutes on a PCR block, then incubated in a humid chamber at 37°C overnight. Slides were washed once in 2X SSC, once in 2X SSC+Hoechst 33342 and once in 2X SSC. Mounting media was added and the slides were imaged.

#### RNA FISH

Slides were prepared for RNA FISH using the same protocol as for DNA FISH but with the RnaseA treatment omitted. Xist RNA FISH was performed using a mixture of Cy3-labeled DNA oligos covering Repeats A, B and C within *Xist*. The RNA FISH protocol was the same as the DNA FISH protocol, except that the denaturing step was omitted and the hybridization buffer+probe mixture was heated at 92°C for 5 minutes then 37°C for 5 minutes and then added directly to the slides. Slides were incubated at 42°C for 4-8 hours and then were washed once in 2X SSC, once in 2X SSC+Hoechst 33342 and once in 2X SSC. Mounting media was added and the slides were imaged.

#### Immunofluoresence

75,000-150,000 cells were cytospun onto slides for 5 minutes at 1000 rpm. Slides were washed once with 1X PBS, then 1X PBS+4% formaldehyde was added for 10 minutes at room temperature, then 1X PBS+0.5% Triton-X 100 for 10 minutes at room temperature to remove un-crosslinked proteins. Slides were washed once in 1X PBS, excess buffer was removed from cell spots and 1% BSA in 1X PBS was added for 45 minutes. Block solution was removed and a 1:200 dilution of H3K27me3 antibody (Active Motif 39155) in 1X PBS+1% BSA was added for 1 hour. Slides were washed 3X in 1X PBS+0.02% Tween-20. Excess liquid was removed and a 1:2000 dilution of goat-Anti-Rabbit Alexa 555 conjugated antibody (ThermoFisher) was added for 1 hour in the dark. Slides were washed once in 1X PBS+0.02% Tween-20, then twice in 1X PBS and then imaged.

#### ImmunoFISH

Slides were prepared the same way as for immunofluorescence; with 0.5 U/uL Protector RNase Inhibitor (Sigma) added to the blocking buffer. To visualize the nucleolus, we used a 1:200 dilution of Nucleophosmin antibody (abcam 10530) in blocking solution as the primary antibody. After immunofluorescence, we post-fixed the slides for 10 minutes in 4% formaldehyde+PBS, and then Xist RNA FISH was performed starting at the dehydration step.

#### Metaphase immunofluorescence

We added 50 ng/mL Karyomax to the media of day 10 differentiating embryoid bodies for 4 hours to arrest cells in metaphase. We harvested the cells via trypsinization, and trypsin was quenched by addition of media. We spun the cells at 1000 rpm for 5 minutes, aspirated the media, then washed twice in 1X PBS. Cells were then resuspended to a concentration of 5×10^5^ cell/mL in 75 mM KCl, and placed at 37°C for 10 minutes for swelling. 1×10^5^ cells were then cytospun onto a microscope slide at 1000 rpm for 5 minutes. The cells were fixed in PFA and immunofluorescence was performed as described for interphase cells. We stained H3K27me3 with a 1:200 dilution of Active Motif 39535 and H3K27ac with a 1:200 dilution of Cell Signaling D5E4.

#### Hi-C library preparation

We used the *in situ* Hi-C method of Rao et al.^3^ to prepare all libraries, using 5-10 million cells. Importantly, we sequenced 20-40 million reads per library. This is a lower sequencing depth than many published Hi-Cs, however since the megadomains are large and prominent feature of the organization of the Xi, this depth is appropriate for detecting the megadomains efficiently and economically. We performed a timecourse of Hi-C experiments at 4 timepoints during differentiation (days 0, 3, 7 & 10). To test whether megadomains form in the absence of *Dxz4* or *Firre*, we differentiated cells for 10 days and performed Hi-C in wild-type, *Dxz4*^∆/∆^, *Firre*^Xi∆/+^ or *Dxz4*^∆/∆^:*Firre*^Xi∆/+^. Finally, to test whether megadomains can form on an autosome with *Xist* and *Dxz4* ectopically inserted, we performed Hi-C in the *Xist+Dxz4* transgene line after 2 days of induction with 1000 ng/mL dox, as well as an *Xist+P* only transgene line after 2 days of induction with 1000 ng/mL dox and the parental male XY rtTA line (no induction).

#### Hi-C analysis

Hi-C alignment to mm9 was performed according to the method of Minajigi & Froberg et al.^4^ The allele-specific Hi-C reads were filtered for quality and uniqueness with HOMER. Custom scripts were used to convert HOMER tag directories into the format accecpted by Juicebox; contact maps were generated using the Juicer tools ‘pre’ command. All Hi-C contact maps visualized in this study are KR-normalized contact maps generated by Juicebox.

The first principal component of the Hi-C correlation matrix has been used as a quantitative measure of the presence or absence of megadomains^5^. We used R to generate the Pearson correlation of 1Mb KR-normalized allele-specific chrX Hi-C matrices, and we plot the first principal component as a function of position along the Xchromosome. Hi-C matrices with a megadomain exhibit a sharp transition in the first principal component score at the bin containing *Dxz4*.

#### Hi-C mixing experiment

We mixed together aligned reads from the day 0 and day 10 Hi-C libraries such that 0%, 10%, 25%, 50%, 75% and 100% of reads were from the day 10 Hi-C. We the generated HOMER tag directories and normalized contact maps in Juicebox as described for the Hi-C experiments. We plotted PC1 scores across the Xi at 1Mb resolution, and defined the PC1 slope at Dxz4 as the PC1 score @ bin 74 – PC1 score @ bin 72.

#### HYbrid Capture Hi-C (Hi-C^2^)

HYbrid Capture Hi-C (Hi-C^2^) probes were designed and hybridization to *in-situ* Hi-C libraries carried out as described previously^6^. Probe sets were designed to enrich interactions in two regions of interest: chrX:70,370,161-71,832,975 and chrX:71,832,976-73,511,687 (mm9). Briefly, 120 bp probes were designed around the MboI restriction sites of the regions of interest as previously described ^6^ and custom synthesized pools of single stranded oligodeoxynucleotides ordered from CustomArray, Inc. (Bothell, WA). Single stranded DNA oligos were amplified and biotinylated in a MAXIScript T7 transcription reaction (Ambion). The resulting biotinylated RNA probes were hybridized to 250-300 ng of *in situ* Hi-C libraries for 24 hours at 65C. DNA hybridized to the RNA probes was pulled down by streptavidin beads (Dynabeads MyOne Streptavidin C1, Life Technologies), washed, and eluted as described ^6^. The resulting DNA was desalted using a 1X SPRI cleanup and amplified with Illumina primers for 18 cycles to prepare for sequencing.

Hi-C^2^ libraries were sequenced to a depth of 8-15 million 50 bp paired-end reads. Reads were trimmed using cutadapt with the options --adapter=GATCGATC (MboI ligation junction) and --minimum-length=20. Reads of each pair were individually mapped to the mus and cas reference genomes using novoalign and merged into Hi-C summary files and filtered using HOMER as previously described ^4^. For the chrX:70,370,161-71,832,975 captures, 3-4% of mapped and paired reads fell within the target region (0.05% expected based on size of capture region versus genome) and for the chrX:71,832,976-73,511,687 captures, 1-2% of mapped and paired reads fell within the target region (0.06% expected based on size of capture region versus genome). To avoid computational complexities arising from normalization of sparse, non-enriched regions in the Hi-C contact map, only Hi-C interactions falling within the capture region were analyzed further. For each capture, a custom script was used to pull out the filtered Hi-C interactions falling within the target region from the HOMER tag directories. Hi-C contact maps of the capture regions were then generated from these HOMER tags using the ‘pre’ command of Juicer tools ^7^. The resulting Hi-C contact maps in .hic format were visualized and normalized with the ‘Coverage (Sqrt)’ option in Juicebox^8^.

#### Insulation score analysis with Hi-C^2^ data

We computed insulation score across the *Mecp2* and *Dxz4* regions to quantitatively measure changes in domain organization during the timecourse of X-inactivation. To do this, we output the ‘Coverage (Sqrt)’ normalized Hi-C contact maps at 25 kb resolution across either the *Mecp2* or *Dxz4* regions using Juicer tools ‘dump’ command. We used custom shell and R scripts to convert the densematrix format output from Juicer into the full matrix format accepted by the cworld suite of Hi-C tools (https://github.com/dekkerlab/cworld-dekker). We computed insulation scores across the captured regions using the cworld perl script ‘matrix2insulation.pl’ using the parameters ‘-v --is 125000 --ids 75000 –im sum’. This set of options uses a smaller number of bins to calculate insulation scores, which we found to be optimal for analyzing insulation over small regions with just a few dozen bins. We plotted the distribution of insulation scores across each region and each timepoint. We evaluated changes in insulation across regions by testing whether there was a difference in the variance of insulation scores between timepoints or between the Xa and the Xi using the F-test. This is appropriate as a loss of insulation by definition is a decrease in the variance of insulation across a region ^9^, which can be visualized as a “flatter” insulation score curve. To generate violin plots and calculate F-test p-values, we excluded the 6 bins on the left and right edges of each Hi-C^2^ region because the windows used to calculate insulation score at these loci fall partially outside the region covered by Hi-C^2^ probes and have far less read coverage than the regions covered by the probes.

#### 4C library preparation and analysis

We previously developed a modified 4C protocol^10^ to examine chromatin conformation from repetitive viewpoints. Our protocol has several advantages over existing 4C profiles: 1.) It sequences the genomic region amplified by the 4C primers, ensuring that on-target priming events can be identified and filtered from numerous off-target priming events 2.) Sequencing the viewpoint allows every read to be assigned to a particular allele if the viewpoint is near a variant, 3.) We use a random barcode to identify PCR duplicates, which previously has not been possible in 4C experiments. We performed our modified 4C using the protocol and analysis pipeline previously described for viewpoints within PAR-TERRA repeats^10^. We used it for several viewpoints within *Firre* and *Dxz4*. Some viewpoints were in the core tandem repeats. For these viewpoints, we use the read outside the viewpoint for allelic determination. Others were in unique regions near the tandem repeats; for these we could use known variants to assign every read to the Xa or the Xi. We performed our analysis in two fibroblast lines, one where the mus X is inactive (mus Xi cas Xa), the other where the cas X is inactive (mus Xa cas Xi).

#### Assay for Transposase-Accessible Chromatin with high-throughput sequencing

50,000 cells were washed in cold PBS and lysed in cold lysis buffer (10 mM Tris-HCl, pH 7.4, 10 mM NaCl, 3 mM MgCl2, 0.1% IGEPAL CA-630) containing proteinase inhibitor cocktail (Roche). Nuclei were resuspended in 1X TD Buffer (Illumina FC-121-1030) and 2.5uL of Tn5 Transposase (Illumina FC-121-1030) were added. Transposition reaction was performed at 37°C for 30min, and DNA was purified using a Qiagen MinElute Kit. DNA libraries were amplified for a total of 8 cycles. Libraries were assessed for quality control on the BioAnalyzer 2100 (Aglient) to ensure nucleosomal phasing and complexity. Sequencing was performed on the HiSeq 2500 (Illumina), using 50 bp paired-end reads.

#### ATAC-seq analysis

Attack seq alignment to mm9 was performed exactly as ChIP-seq alignment was performed in Minajigi & Froberg et al. ^4^. Peaks were called using macs2 with default parameters. Biallelic peaks were identified as peaks with at least 10 alleleic reads in a sample and an Xi:Xa ratio greater than 1/3. Xi-specific peaks were defined as peaks with at least 10 allelic reads and a Xi:Xa ratio less than 1/3. To test whether Xi-specific peaks in wild-type are “restored” (that is: acquire appreciable accessibility on the Xi) in either the *Dxz4* or *Firre* deletion, we plot the wild-type Xa reads on the x-axis and the deletion Xi reads on the y-axis and identify peaks where the deletion Xi/wild-type Xa ratio is greater than ½ (these are peaks where the deletion accessibility level reaches at least half the wild-type accessibility ratio). We also examine the biallelic peaks and plot the wild-type Xi reads on the x-axis and the deletion Xi reads on the y-axis to determine whether the accessibility on the Xi changes for the peaks that are bi-allelic in wild-type.

#### RNA-seq library preparation

Total RNA was isolated from 2-5 million trypsinized cells using trizol extraction. polyA+ mRNA was isolated using the NEBNext NEBNext® Poly(A) mRNA Magnetic Isolation Module using 5 ug of total RNA as input. Isolated mRNA was reverse-transcribed using Superscript III and actinomycin D to inhibit template switching. Second-strand synthesis was performed using the NEBNext Ultra Directional RNA Second Strand Synthesis Module. Library preparation and NEBNext® ChIP-Seq Library Prep Master Mix Set for Illumina. A USER enzyme treatment was performed following adaptor ligation to specifically degrade the second strand and allow a stranded analysis. Libraries were amplified for 10-15 cycles of PCR using Q5 polymerase and NEBNext multiplex oligos.

#### RNA-seq analysis

RNA-seq reads were aligned to the cas (Xa) and mus (Xi) genomes allele-specifically using a previously published pipeline ^4,11-13^. Following alignment, gene expression levels for each gene were defined using HOMER. Differential expression and fold changes between conditions were calculated using DESeq2. We plotted the cumulative distributions of fold changes for autosomal and X-linked genes and evaluated the significance of any differences between the distributions of the fold changes using the Kolmogorov-Smirnov (KS) test. To examine allele-specific expression from the Xa and the Xi, we summed together allelic reads across both biological replicates and filtered for genes with at least 12 allelic reads in both wild-type and *Dxz4*^∆/∆^:*Firre*^Xi∆/+^and fpm > 0 in all replicates. We also used RNA-seq done in pure hybrid mus or cas fibroblasts to identify and eliminate genes that have incorrect SNP information. We defined escapee genes in a particular condition as genes where at least 10% of allelic reads came from the Xi in either replicate of that condition. We plotted the distribution of expression levels from the Xi (Xi/(Xi+Xa) read counts) for all genes passing our filtered for each replicate. We evaluated the significance in differences of the mean expression level from the Xi using the Wilcoxon Signed Rank Test with Bonferroni correction for multiple hypothesis testing.

#### qRT-PCR

Total RNA was isolated from cells using trizol extraction. 500 ng RNA was heated at 70 degrees C for 10 minutes then cooled to 4 degrees in the presence of 50 ng random primers in 5 uL total volume. The RNA was reverse-transcribed in a 10 uL reaction containing 1X First Strand buffer, 10 mM DTT, 500 uM dNTPs, 6U Protector RNase inhibitor and 100U Superscript III. The reaction was incubated for 5 minutes at 25 degrees, then 1 hr at 50 degrees and 15 minutes at 85 degrees. Reverse transcription reactions were diluted to 100 uL with water before qPCR. 500 ng RNA was added to 100 uL water as a -RT control. 1 uL template was used per 15 uL qPCR reaction prepared with 1X Taq UniverSYBR Green (BioRad) master mix and 200 nM primers, and reactions were performed in triplicate. All qPCR primers were run using an annealing temperature of 55 degrees.

